# Functional imaging of 3D bioprinted microalgal constructs and simulation of their photosynthetic performance

**DOI:** 10.1101/2025.02.19.639067

**Authors:** Swathi Murthy, Maria Mosshammer, Erik Trampe, Michael Kühl

**Affiliations:** Marine Biological Section, Department of Biology, University of Copenhagen, Strandpromenaden 5, DK 3000 Helsingør, Denmark

## Abstract

The intricate three dimensional architecture at different spatial length scales affects the functionality and growth performance of immobilized photosynthesizing cells in biofilms and bioprinted constructs. Despite the tremendous potential of 3D bioprinting in precisely defining sample heterogeneity and composition in spatial context, cell metabolism is mostly measured in media surrounding the constructs or by destructive sample analyses. The exploration and application of non-invasive techniques for monitoring physico-chemical microenvironments, growth and metabolic activity of cells in 3D printed constructs is thus in strong demand. Here, we present a pipeline for the fabrication of 3D bioprinted microalgal constructs with a functionalized gelatin methacryloyl (GelMA)-based bioink for imaging O_2_ dynamics within bioprinted constructs, as well as their characterization using various, non-invasive functional imaging techniques and numerical simulation of their photophysiological performance. This fabrication, imaging and simulation pipeline now enables investigation of the effect of structure and composition on photosynthetic efficiency of bioprinted constructs with microalgae or cyanobacteria. It can facilitate designing efficient construct geometries for enhanced light penetration and improved mass transfer of nutrients, CO_2_ or O_2_ between the 3D printed construct and the surrounding medium, thereby providing a mechanistic basis for the design of more efficient artificial photosynthetic systems.

## 1. Introduction

Functionality and growth performance of photosynthesizing cells organized in biofilms, tissues of organisms or free-living, can be linked to the intricate three-dimensional architecture at different spatial scales ranging from canopies in forests^1^ to small scale modulation of scattering and absorption e.g. in the leaves of terrestrial plants^2^, aquatic biofilms^3 4^ and individual microalgae^5,6^. For example, the high photosynthetic efficiency of symbiotic algae in corals can be attributed to efficient light management in coral holobionts arising from their complex shapes and optical properties.^7,8^ Coral studies have inspired the development of efficient light harvesting structures^9^, which can alleviate the limited light penetration in natural phototrophic biofilms^10^ and hence improve their photosynthetic efficiency and potential for biofuel production.^11–15^ Such bioinspired materials studies take advantage of the rapid advancement in 3D additive manufacturing technologies combined with bio-compatible materials, which have enabled precise 3D patterning of biomaterials (hydrogels) containing living cells, using automated and cell-friendly printing processes known as 3D bioprinting.^16–19^ Application of 3D bioprinting allows for fast prototyping and up-scaling of design solutions for large scale implementations, and this technique is extensively applied for tissue engineering and organ printing studies, as well as for encapsulating human and other mammalian cells in hydrogels.^19–23^ In comparison, relatively few studies have investigated the potential of 3D bioprinting for other biotechnological applications^24,25^ using e.g. microbes such as bacteria^26–28^ or photosynthetic microalgae^9,29–31^ for production of biofuels and other biomaterials.^32^ Microorganisms can produce a wide range of secondary metabolites that can be used in food, medicine, cosmetics, pharmaceuticals and energy industry.^33,34^ However, their small size can be a drawback due to the requirement of additional cell separation processes.^35^ Currently, microalgal harvesting requires a rigorous step of water removal by methods, such as centrifugation, filtration, electrical-based processes and flotation.^36^ Here immobilization techniques like bio printing can facilitate high density cultivation^37^, along with easy and efficient biomass harvesting^38^.

The use of 3D bioprinting in ecology and ecophysiology is still rare despite its tremendous potential for i) preparation of samples with defined heterogeneity and composition, and ii) the targeted investigation and modulation of specific traits in a spatial context. Wangpraseurt et al.^9,39^ e.g. investigated the possibility of 3D printing bionic analogues to living corals to learn more about their outstanding light management systems and to develop bioinspired 3D printed constructs for efficient microalgal growth. The possibility to control composition and biomass distribution in bioprinted constructs has e.g. been used in several studies that combined 3D bioprinting of algae and cyanobacteria with animal cells^31^, as a way to overcome the inherent O_2_ limitation in larger bioprinted constructs. Such O_2_ limitation remains a major issue^40,41^, e.g. in tissue engineering applications, due to the lack of a vascular network in bioprinted constructs and a lack of suitable techniques for dynamic assessment of how construct design affects O_2_ distribution and availability. In a proof-of-concept study, Trampe et al. (2018) developed first bioinks functionalized with O_2_-sensitive luminescent nanoparticles and used 3D bioprinting to deposit sensor particles, living algae and stem cells in an alginate-based hydrogel scaffold^9,29–31^. This novel approach enabled first non-invasive real-time monitoring of spatio-temporal O_2_ dynamics in bioprinted cell constructs, and such combination of optical O_2_ sensing and bioprinting has many applications e.g. in advanced tissue constructs^42^.

With increasing popularity of 3D bioprinting, the exploration and application of non-invasive techniques for monitoring physico-chemical microenvironments, growth and metabolic activity of cells in 3D printed constructs is in strong demand.^16,24,43^ Many potential techniques have been successfully used to study natural systems like biofilms, animal tissue and aquatic/terrestrial organisms. For example, optical O_2_ sensing in aquatic systems^44^, micro-sensor measurements of O_2_, H_2_S and other chemical species in natural environments^45^, light sheet microscopy on live biological specimen^46^, optical coherence tomography (OCT) on intact corals and in dermatology^47–49^, or hyperspectral imaging of the rhizosphere of aquatic plants^50^. Applying these techniques to monitor structure, microenvironments and cell function relationships in bioprinted constructs, combined with numerical simulations of light and the chemical microenvironment^51^ can help with design optimization and to study biological phenomena relevant for specific biotechnological and pharmaceutical applications.

In this study, we present a pipeline for the fabrication of 3D bioprinted, microalgal constructs with a functionalized gelatin methacryloyl (GelMA)-based bioink for imaging O_2_ dynamics within the prints, as well as their characterization using various, non-invasive functional imaging techniques and numerical simulation of their photo-physiological performance. The light distribution in the print was characterized by measuring the directly back scattered light from the samples by optical coherence tomography (OCT)^49^. Pulse-amplitude-modulated, variable chlorophyll fluorescence imaging (VCFI) was used to estimate the health of the micro-algae^52^. Photosynthesis and respiration in the bioprinted constructs were quantified in novel experimental chambers enabling simultaneous determination of i) total net O_2_ consumption/production via gas exchange measurements, and ii) imaging of local O_2_ dynamics over the bioprinted constructs via luminescence lifetime imaging.^53,54^ Simulation of the photo-physiological performance of the prints was carried out by coupling finite element Monte-Carlo (MC) simulation of radiative transfer^55^ with diffusion-reaction modeling of photosynthesis, respiration and mass transfer of O_2_ between the bio-print and the surrounding medium.^51^ This fabrication, imaging and simulation pipeline now enables investigation of the effect of structure and composition on photosynthetic efficiency of bioprinted constructs with microalgae or cyanobacteria. It can facilitate designing efficient (surface) topographies for enhanced light penetration and improved mass transfer of nutrients, CO_2_ or O_2_ between the 3D printed construct and the surrounding medium thereby providing a mechanistic basis for the design of more efficient artificial photosynthetic systems^56–60^.

## 2. Materials and methods

Gelatin from porcine skin (gel strength 300, Type A), methacryl anhydride (containing 2,000 ppm topanol A as inhibitor, 94%), NaHCO_3_, and poly(1-vinylpyrrolidone-co-styrene) (38% emulsion in H_2_O, <0.5 μm particle size) were purchased from Sigma Aldrich (sigmaaldrich.com) and used without further modification. The tangential flow filtration (TFF) system was generously provided by PALL (pall.com/), and the MinimateTM TFF capsule with a 10K omega membrane was purchased from PALL. Platinum(II)meso-tetra(4-fluorophenyl)tetrabenzoporphyrin (PtTPTBPF) was generously supplied by Dr. Sergey Borisov (Institute for Analytical Chemistry and Food Chemistry, Graz University of Technology (tugraz.at/institute/acfc/home/). Macrolex® fluorescence red was bought from Kremer Pigments (shop.kremerpigments.com/en/). TAP medium was prepared in MilliQ water, according to the recipe: https://utex.org/products/tap-medium?variant=30991736897626#recipe. 0.25% solution of Trypsin-EDTA was purchased from Sigma Aldrich (sigmaaldrich.com). Irgacure 2959 was purchased from BASF (IC2959; BASF, cat. no. 029891301PS04) and Lithium phenyl-2,4,6-trimethylbenzoylphosphinate from Sigma Aldrich (LAP; Sigma Aldrich)

### 2.1 GelMA synthesis

Two gelatin methacryloyl (GelMA)–based hydrogels with different degrees of functionalization (DoF) were synthesized according to literature^61^. In short, 10 g gelatin were swelled for 30 minutes at room temperature and fully dissolved at 50°C in a water bath under stirring in 100 mL MilliQ water. 4 g (DoF_1_) or 6 g (DoF_2_) methacrylic anhydride were added dropwise to the vigorously stirred gelatin solution. After addition of methacrylic anhydride, all further steps were conducted at low light levels or darkness. The solution was stirred for 1h and subsequently centrifuged at 3500 rpm for 3 minutes at room temperature. The pellet was discarded and the clear supernatant diluted with 2 volumes deionized water heated to 40 °C. The diluted solution was then dialyzed at 40°C for 5 days using a Minimate^TM^ TFF capsule (10 kDa cut-off). The pH was adjusted to 7.4 using NaHCO_3_ (1M), and the solution was filter-sterilized using a vacuum filtration unit with a 0.2 µm filter (PES membrane). The GelMA solution was transferred to 50 mL vials with vented screw caps, snap frozen and lyophilized at −55°C in a freeze dryer (CoolSafe 55-4 Pro, Ninolab) for 7 days. The lyophilized GelMA was stored at −20°C prior to usage.

### 2.2 Algal culture

A culture of the green alga *Chlorella sorokiniana* UTEX1230 was maintained in culture tubes with TAP medium (see above) under constant shaking and under a defined photon irradiance from white LED’s (10 μmol photons m^−2^ s^−1^; 400-700 nm) in a temperature-regulated culture cabinet (ALGAETRON, Photon System Instruments) at 24⁰C.

### 2.3 Optical sensor nanoparticle synthesis

The O_2_-sensitive nanoparticles (NP) were prepared following literature procedures for staining poly(1-vinylpyrrolidone-co-styrene) (PS-PVP) beads with an average particle diameter of ∼ 116 nm^62,63,64^ In short, 526 mg of PS-PVP were diluted in a mixture of 50 mL MilliQ water and 30 mL tetrahydrofuran (THF). 3 mg of the O_2_-sensitive indicator dye platinum(II)meso-tetra(4-fluorophenyl)tetrabenzoporphyrin (PtTPTBPF) and 3 mg of the antenna dye Macrolex® fluorescence red were dissolved in 20 mL THF and added dropwise to the vigorously stirred PS-PVP emulsion. THF was removed by blowing an airstream over the surface and the volume of the suspension was reduced at 50°C to a final concentration of ∼ 6.5 mg particles mL^-1^.

### 2.4 Functionalized bio-ink preparation

A photo-initiator stock solution was prepared by dissolving 250 mg of Irgacure 2959 or Lithium phenyl-2,4,6-trimethylbenzoylphosphinate (LAP) in 10 mL TAP medium at 70⁰C with constant stirring and allowed to cool to room temperature.^61^ 8% (w/v) GelMA ink was prepared, by soaking GelMA foam in TAP medium with photo initiator (to final concentrations, w/v, of 1% for Irgacure and 0.025% for LAP) overnight in the refrigerator. This was followed by dissolving the GelMA in an incubator at 37⁰C while stirring at 300 rpm until the mixture turned clear.^61^ The pH of the solution was adjusted to pH 8.1 by adding a strong base (1M NaOH). The GelMA ink was finally autoclaved at 80⁰C for 15 minutes to ensure its sterility. The ink was cooled to <35 ⁰C, before adding microalgal cells and the sensor NP solution. The NP sensor solution (10% volume fraction) was mixed with the bioink. The algal cells were spun down from a concentrated stock solution at 9000 rpm. The supernatant was discarded and the algal cells were added at a concentration of ∼10^8^ cells mL^-1^ bioink for high biomass load and 20 x 10^6^ cells mL^-1^ bioink for low biomass load.

### 2.5 Bioprinting

Two different GelMA inks with varying degree of functionalization were tested for bioprinting. While both of them exhibited good viability for algal growth, the ink with higher degree of functionalization showed better long term stability of the bioprinted constructs. Two simple designs, i.e., a square slab and a construct with v-grooves (60⁰ vertex angle and 2 mm height)^15^ of same total volume (57.6 mm^3^) and foot print (48 mm^2^), were bioprinted. The constructs were designed in a CAD software (Autodesk Fusion 360.ink), followed by slicing and g-code creation in PrusaSlicer (PrusaSlicer.ink). The g-code was uploaded and printed on a commercial 3D bioprinter (BioX, CellInk Lifesciences), which utilizes an extrusion based printing technique. The printing was done with an infill density set to 60% using a 3D honeycomb pattern. The print speed and extrusion pressure were set to 1 mm s^-1^ and 25 kPa, respectively. The temperature of the bio-ink was adjusted to 24⁰C, to obtain a desired viscosity for printing. The constructs were printed, on a PET foil substrate maintained at 10⁰C and allowed to stand for 5 minutes (after printing the entire construct) to facilitate physical cross-linking, before curing with 365 nm or 405 nm light. The UV light (365 nm) curing was done at a light intensity of 2 mWcm^-2^ for 7 minutes, whereas 405 nm curing was done light intensity of 3 mWcm^-2^ for 30 seconds. The prints were stored in sterile petri dishes with TAP medium sealed off at the edges with parafilm. Printing and handling was carried out in a sterile environment, either in the bioprinter (fitted with UV-C germicidal lamps and a HEPA H14 dual-filter system) or in a laminar flow bench. Prior to and between experiments, the prints were stored in an incubator (AlgaeTron AG 230) under constant low photon irradiance (400-700 nm) of 10 μmol photons m^−2^ s^−1^ from white LED’s

### 2.6 Cell counts

The cell density was determined at the beginning of the experiments (day 0) by measuring the cell concentration in the algal stock solution, before adding it to the bio-ink. Cell concentration in the bioprints during the growth phase was determined by removing the prints from the growth medium and dissolving them in 600 µL trypsin solution^65^ (0.25% Trypsin/EDTA) at 37⁰C for 1 hour. For cell counting, the algal solution (stock culture or dissolved prints in trypsin) was diluted 500-1000 times, depending on the concentration, in TAP medium. 1 mL of the diluted algal solution was poured into a Sedgewick rafter counting chamber, which subdivides 1 mL into (1000) 1µL volume fractions. The number of algal cells in individual 1 µL volume fractions was manually counted under an optical microscope. At least 15 - 20 volume fractions were counted for each sample and averaged (Figure S7).

### 2.7 Optical coherence tomography

A spectral domain OCT system (Ganymed II; Thorlabs GmbH, Dachau, Germany) equipped with an objective lens with an effective focal length of 18 mm and a working distance of 7.5 mm (LSM02-BB; Thorlabs GmbH, Dachau, Germany) was used for OCT imaging.^66^ The system is equipped with a 930 nm light source, yielding a maximal axial and lateral resolution in water of 5.8 μm and 8 μm, respectively. Two dimensional OCT b-scans were acquired at a fixed pixel size of 581 x 1023. The actual field of view was variable in y but fixed in z (= 2.2 mm). The OCT system was optimized to yield highest signal at a fixed distance, in the upper 1/3^rd^ of the image.^66^ OCT imaging was performed on bio-printed constructs fully immersed in TAP medium in a Petri-dish.

OCT system calibration and optical parameter extraction were performed according to published procedures.^67–69^ Briefly, OCT signal calibration was performed before measurements on the bioprints, by using a reflectance standard composed of air-glass, water-glass and oil-glass interfaces. The reflectivity (R) was calibrated via Fresnel’s equation, R = ((n_1_-n_2_)/(n_1_+n_2_))^2^, using n of air (1), water (1.33), oil (1.46) and glass (1.52).^67^ The OCT signal from the measurements on the bioprints (in dB) was converted to R values using a linear fit of log_10_(R) versus OCT intensity values in dB (Figure S11). The OCT scans were also corrected for the focus function of the objective lens via calibration measurements performed in steps of 0.1 mm from z = 0.4 mm (focal plane) to z = 0.8 mm. The OCT signal fall-off from the focal plane follows an exponential decay function. The acquired OCT scan (of the samples) was corrected by dividing with the exponential fit function. Corrected OCT reflectivity values were plotted against sample depth (z, distance from focal volume) and fitted to the exponential decay function (Eq. 1):

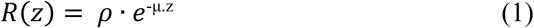

where *ρ* (dimensionless) is the light intensity and µ is the signal attenuation coefficient (cm^-1^) from the focal volume.^67^

Using the grid method^70^, values of *ρ* and µ were matched to estimate the anisotropy of scattering (g) and scattering coefficient (µ_s_; cm^-1^) based on the theory described in the literature.^67,69^ For this, it was assumed that the absorption coefficient, µ_a_, was dominated by water (µ_a_ ∼ 0.43 cm^-1^), while absorption by algae at 930 nm was negligible. The refractive index of the bioprint was assumed to be 1.33, as the surface of a clear GelMA bioprint (without algae) immersed in water was hard to identify under the OCT. The effective numerical aperture (NA) of the objective was 0.11. Curve fitting was performed on randomly selected spots in the bioprinted construct. The fit was considered satisfactory if the model matched the data with a R^2^ value >0.5.^67^

### 2.8 Variable chlorophyll fluorescence imaging

Variable chlorophyll fluorescence (VCFI) is a widely applied non-invasive technique to assess the photosynthetic performance of plants and algae. It relies on externally applied, weak and non-actinic light pulses quantifying the fluorescence of photosynthetic organisms with minimal sample manipulation.^52,71,72^ Changes in the variable Chl *a* fluorescence in response to actinic light and strong saturation pulses are measured to quantify the fate of the light energy absorbed by the photosynthetic apparatus and the balance between photochemical and non-photochemical quenching.^52^ This indirectly informs about the redox status of the photosystem II (PS II). In this study, VCFI was used as a quick qualitative reference to estimate the health of the microalgae and estimate saturating light levels for algal photosynthesis.

The photosynthetic activity of the microalgae in the bioprinted constructs was measured via a pulse-amplitude-modulated, variable chlorophyll fluorescence imaging system (I-PAM/GFP, Walz GmbH, Effeltrich, Germany).^52,72^ The system utilizes blue (470 nm) LED light for weak (<1 μmol photons m^−2^ s^−1^) modulated measuring light pulses, strong (0.8 s at >2500 μmol photons m^−2^ s^−1^) saturating light pulses, and defined levels of actinic irradiance, as measured with a calibrated photon irradiance meter at the level of the bioprinted constructs (ULM, Walz, Effeltrich, Germany).

From measurements of the minimum fluorescence yield, *F_0_*, and the maximum fluorescence yield, *F_m_*, in dark acclimated samples recorded before and during a saturation pulse, respectively, we calculated the maximum quantum yield of PSII as *F_v_/F_m_ = (F_m_-F_0_)/F_m_*. This parameter is frequently used to indicate the health and photosynthetic capacity of photosynthetic organisms^73^. The effective PS II quantum yield in light-exposed samples was calculated as *YII = F^’^_m_-F)/F^’^_m_*, where *F’_m_* is the maximal fluorescence yield of light acclimated samples (under the saturating light pulse) and *F* is the fluorescence yield under ambient actinic light conditions.^52,71^

From YII measurements over a range of increasing actinic photon irradiance levels of photosynthetic active radiation (PAR; 400-700 nm), we calculated a so-called rapid light curve^74^. The RLC quantifies how the relative electron transport rate (rETR = YII x PAR) via PSII changes with increasing photon irradiance and can be used as a qualitative measure of the photo-physiological acclimation state of the micro-algae embedded in the prints.^71^

### 2.9 Monte Carlo simulation of radiative transfer

Radiative transfer in bioprinted constructs kept in a laminar flow chamber was modeled by Monte Carlo (MC) simulations providing a numerical solution to the radiative transfer equation (RTE).^75^ We used a freely available, finite-element voxel-based MC simulation software, i. e., ValoMC^55^, which is based on the algorithm described by Prahl *et al*.^76^. We used MC modelling in combination with COMSOL Multiphysics (v5.6, COMSOL Inc., Burlington, MA) for simulating and visualizing the light field, as well as mass and heat transfer in bioprinted constructs. A detailed description of our approach can be found in our earlier work on multiphysics simulation of radiative, heat and mass transfer in corals.^51^ Briefly, the discretized three-dimensional geometry of the bioprinted constructs (created in COMSOL, with a minimum tetrahedral element size of 3 µm) was loaded into ValoMC. All internal boundaries were subsequently removed. A set of material optical properties was assigned to each of the model domains (summarized in Table S1), consisting of the absorption coefficient *μ_a_*, the scattering coefficient *μ_s_*, the directionality of scattering *g* (using the Henyey-Greenstein approximation for the scattering phase function), and the refractive index *n*. A ‘direct’ light source at 636 nm, launching photons into the flow chamber perpendicular to the upper surface of the construct, was assigned over the entire top boundary with a total power of 1 W. For each simulation, 10^8^ photon packets were launched. The output scalar irradiance from the MC simulation was normalized to the incident photon irradiance. The normalized light field was then imported into COMSOL and mapped over the 3D bioprinted construct by solving the Poisson’s diffusion equation (2) in weak form. The variations in the scalar irradiance data inherent to the stochastic nature of the MC simulation was smoothened using equation (2).

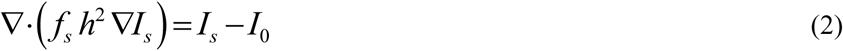

where *I_s_* is the smoothened normalised scalar irradiance, *I_0_* is the imported normalised scalar irradiance, *h* is the local mesh element size, and *f_s_* is a smoothing factor (set to 0.1). A zero light flux condition was applied to all boundaries. Finally, the photon scalar irradiance *I_p_* was calculated from *I_s_*, as shown in equation (3).

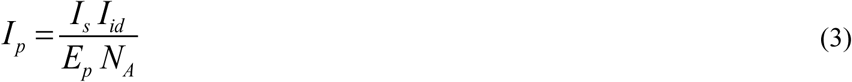

where *I_id_* denotes the incident downwelling irradiance, *N_A_* the Avogadro’s number, and *E_p_* the energy of a photon at 636 nm (λ), calculated as *h*c/λ, *h* is Planck’s constant and *c* is the velocity of light in vacuum).

### 4.10 Mass transfer simulation

Mass transfer simulations were carried out in COMSOL Multiphysics (v5.6, COMSOL Inc., Burlington, MA) using a defined flow chamber geometry.

#### Fluid flow

Stationary and incompressible Navier-Stokes equations for laminar flow were used to simulate the water flow around the bioprinted constructs in a flow chamber (Figure S17):

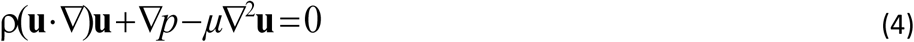

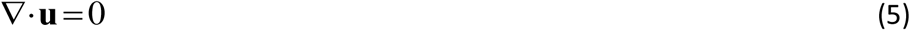

where u is the velocity vector, *p* the pressure, *µ* the dynamic viscosity and ρ the density of water. The water inflow had an average velocity of 5 mm s^-1^, while a zero gauge pressure was set in the outflow. The bottom wall of the flow cell was no-slip (zero-velocity) and symmetry boundary condition was assumed for top and lateral walls of the flow cell. The water velocity profile calculated over the bioprinted construct supported the convective transport of the O_2_, produced due to the outcome of photosynthesis and respiration in the algal constructs.

#### Oxygen transport and reactions

The dissolved O_2_ concentration (*c_O2_*), was calculated from a diffusion-reaction equation (equation 6) in the bioprinted construct and from diffusion-convection equation in the surrounding water column (equation 7):

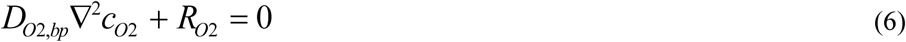

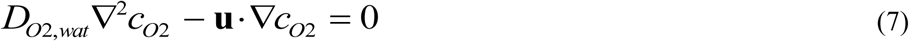

where *R_O2_* is the oxygen reaction rate (net oxygen production) in the bio-print, whereas *D_O2,wat_* and *D_O2,bp_* denote the O_2_ diffusion coefficient in the water and bioprinted construct, respectively. A constant O_2_ concentration (0.25 mol m^-3^) was introduced at the inlet and a convection-only condition at the outlet of the flow cell. All other boundaries were insulated (no flux of O_2_).

The net rate of O_2_ produced by photosynthesis in the bio-print was calculated as:

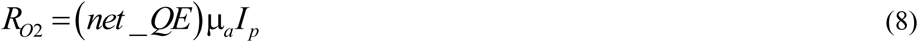

where net_*QE* is the quantum efficiency of algal net-photosynthesis (here assumed to be 0.005 based on QE measurements in natural biofilms^77^), where µ_a_ is the absorption coefficient in the bioprinted construct, and *I_p_* is the photon scalar irradiance. The different parameters used in the simulations are summarized in Table S2.

The flow chamber dimensions (as described in Figure S17) were chosen to reduce both the computational cost and errors in MC, flow and mass transfer calculations. A comparison of simulated light field, for a bioprinted slab geometry (optical properties from day 8, Table S1 and S10), was made between large and small flow chamber scenarios. The results (Figure S14) show that the calculated light field in the bioprint is very similar for both the scenarios. A mesh-independent study (Figure S15) showed that the chosen mesh size, minimum element size of 3 µm for light and 66 µm for O_2_ simulation, is sufficient.

### 2.10 Approximation of optical properties for simulation

The optical properties of bio-prints used for MC simulation were either approximated from the cell count or the OCT data (as described in section 4.7). The algal cell count was used to calculate the optical properties, i.e., the absorption coefficient, µ_a_, and scattering coefficient, µ_s_, based on an earlier experimental study with *Chlorella vulgaris*^78^, a green algae of the same genus as *Chlorella sorokiniana* and with similar size and optical properties, from equations

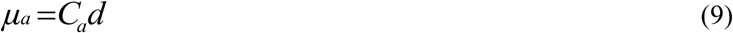

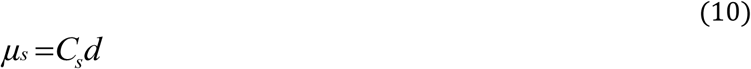

where *C_a_* is the absorption cross-section and *C_s_* is the scattering cross-section at 636 nm expressed in m^2^ kg^-1^, d is algal density expressed in kg m^-3^. Here d was calculated as: (the number of algal cells x dry weight of each cell)/volume of the print. The dry weight of individual algal cells was taken to be 3.9×10^-14^ kg^79^.

The anisotropy of scattering, g, was assumed to be 0.99^78^ and the refractive index, n, for the bioprint was set to be 1.33, as a clear GelMA slab in water was practically invisible under OCT. To account for the algal growth dynamics, OCT images (section 2.3) were used to extract the optical properties µ_s_ and the anisotropy of scattering, g (see section 4.7), for samples on day 8. It was assumed that the extracted µ_s_ at 930 nm would be similar to the value at 636 nm (used for simulations), and µ_a_ was obtained from the number of number algal cells (eq. 9).

### 2.11 Algal growth measurements

Four identical samples of Slab_NP+alg, from the same bio-ink formulation, were printed consecutively and stored individually in a petri dish with TAP medium, in the incubator (as described in 4.5). The slabs were photographed right after printing and one of the slabs (#1) was photographed, every 2 days, during the 8 days of exponential growth phase. One of the slabs was chosen for analysis, each on day 2, 4, 6 and 8. First, VCFI measurements were made on the sample (as described in 4.8), followed by sacrificial cell count measurements (as described in 4.6).

### 2.12 Luminescence lifetime imaging of dissolved O_2_ concentration and respirometry measurements

The O_2_ dynamics in bioprinted constructs with a slab and v-groove configuration was measured with samples kept in a customized, gas-tight glass chamber (10 mL volume) with a flat, glass coverslip as cover, through which imaging was done. The edges of the chamber were painted black, to avoid light guiding and scattering effects. The construct was placed on a small, flat holder inside the chamber above a stirring bar, ensuring constant flow and efficient mass transfer between construct and the surrounding water (Figure S5). An optical O_2_ sensor spot with an optical isolation (PyroScience GmbH) was mounted inside the chamber and could be read out via an optical fiber attached to the chamber at one end and to a fiber-optic O_2_ meter (FireStingO_2_; PyroScience GmbH) at the other end. Calibration of the sensor spots were done according to the manufacturer, and O_2_ concentrations inside the experimental chamber were logged every 6 s enabling a quantification of the net O_2_ exchange between the bioprinted constructs and the surrounding water in the closed chamber.

The net O_2_ exchange (Net rate), where positive values indicate net production and negative values indicate net consumption of O_2_, was calculated from the measured linear change in O_2_ concentration in the chamber over time (Slope) corrected for the water volume surrounding the print (V) and the printed construct surface area (PS):

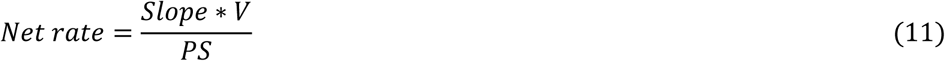

The O_2_-dependent change in the luminescence lifetime of optical sensor nanoparticles in the functionalized bioink was measured over time on constructs using a time-domain luminescence lifetime camera system (PCO Sensicam Sensimod; Excelitas) according to Holst et al. (1998). The camera was equipped with a color-corrected objective (Xenoplan XNP 1.4/23 CCTV-Lens; 400–1000 nm; schneiderkreuznach.com). A 565 nm long-pass filter (041 Rotorange 4X; schneiderkreuznach.com) was mounted in front of the objective. The luminescent sensor particles were excited using a 460 nm LED, triggered by a custom made trigger unit.^80^ Image acquisition and trigger control was done using the dedicated software look@molli v1.82a. For calibration, a clear GelMA slab was printed on a planar optode containing the same dyes as the nanoparticles. Oxygen concentrations were varied by flushing with N_2_. Two calibration curves were determined; directly on the planar optode, as well as through the GelMA slab. No interference by GelMA was observed. As a control, a two point calibration of the functionalized GelMA ink with sensor nanoparticles (air saturated and anoxic conditions) was conducted, which correlated well to the calibration curves using the planar optode. The calibration curves were represented as Stern-Volmer plots and fitted using the two-site model^81^.

For recording of O_2_ dynamics, the bioprinted constructs were kept in darkness for 120 minutes (slab) and 50 minutes (v-groove) prior to switching on the light source, i.e., a fiber-optic tungsten-halogen lamp (K2500 LCD, Schott GmbH) equipped with a fiber optical ring light, which was mounted around the objective of the camera. The incident photon irradiance of photosynthetically active radiation (PAR; 400-700 nm) at the level of the construct surface in the chamber was measured with a calibrated spectroradiometer (BTS 256, Gigahertz Optics GmbH) for different lamp settings, i.e., 115, 260, 356, 450, and 650 µmol photons m^-2^ s^-1^. Lifetime images were recorded either during dark incubation or during light incubations at known photon irradiance by intermittent darkening during image acquisition. The net production of O_2_ as a function of incident photon irradiance was fitted to an exponential function according to Spilling et al.^20^ and the estimated gross photosynthesis according to Webb et al.^21^

## 3. Results and discussion

In this study, we demonstrate a pipeline for imaging the structure, function and metabolic performance of 3D printed hydrogel constructs containing the green microalga *Chlorella sorokiniana* UTEX1230 in two different topographies, i.e., a simple slab and a Vgroove geometry with the same volume and areal footprint. The bioprints were investigated using VCFI, OCT, growth measurements, and measurements of surface dynamics of dissolved O_2_ and bulk water respirometry, supplemented with light and O_2_ mass transfer simulations (Figure 1).

**Figure 1:**
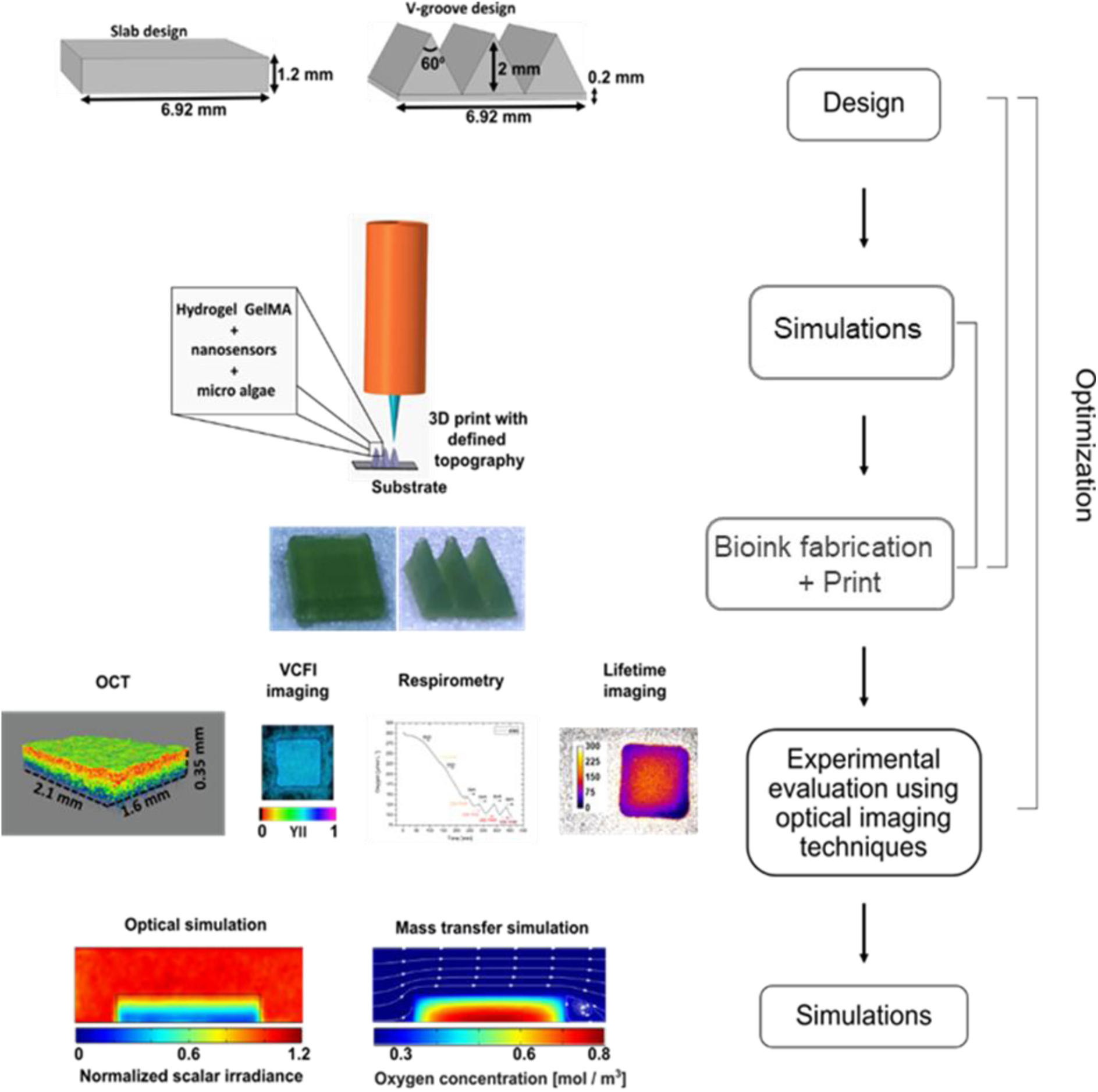
Schematic representation of the pipeline for 3D bioprinting, optical imaging and simulation of micro-algal constructs. **Step 1,** Design: Print layouts are designed and optimized for certain parameters. Step 2, Bioink fabrication: investigating the mixing step, temperature, choice of sensor particles etc. Step 3, Print; different bioinks and their composition (hydrogel : cell amount : optical sensor nanoparticles) are tested for print fidelity, long term stability, etc. and optimized for the print layout. Step 4, Non-invasive experimental evaluation of construct structure, with optical coherence tomography (OCT), and function using variable chlorophyll fluorescence imaging (VCFI), and measurements of O_2_ concentrations using respirometry and lifetime imaging of O_2_ sensor particles for quantifying photosynthesis and respiration. Step 5, 2D and 3D simulations of normalized scalar irradiance and the O_2_ distribution for a given incident irradiance within and around a bioprinted construct in a laminar flow chamber.

The samples were printed using a functionalized gelatin methacryloyl (GelMA)-based bioink, containing O_2_ sensitive optical sensor nanoparticles (NPs)^53,54,82^. GelMA was chosen due to its high optical clarity to minimize the potential interference of the material with the optical imaging techniques.^61^ The rheological properties of GelMA can be easily tuned with slight modification in temperature^83^ (within the tolerance limits of the used algal cells), yielding good control over its print fidelity. The bioprinted constructs were made with sterile inks and sterile work procedures and showed long term stability and algal growth for at least 5 weeks after printing, provided they were stored under low light conditions. Non-sterile procedures resulted in degradation of the prints within 7 to 8 days (Figure S1) and poor algal growth, due to accelerating bacterial growth and breakdown of the cross-linked GelMA hydrogel matrix.

Two different types of photoinitiators, cross-linking at either 360 nm (Irgacure 2959) or 405 nm (Lithium phenyl-2,4,6-trimethylbenzoylphosphinate; LAP) were investigated with varying algal biomass loads in the bioink. Using UV-curing, we observed bleaching and growth inhibition of the microalgae in the outer layers of the bioprinted constructs, especially with a high biomass load, as this required a higher dose of UV radiation for curing, which is also absorbed by algal DNA^84^ and can cause photo-bleaching (Figure S2). Algal cell viability also declined with increasing curing dose and increasing LAP concentration. Highest viability of algae were obtained by using LAP in combination with a low initial biomass load (20 x 10^6^ cells mL^-1^) of green microalgae in the bioink showing best algal cell viability (Figure S3). The optimized curing time for 405 nm light at 3 mWcm^-2^ was 30 seconds, using a LAP concentration (w/v) around 0.025%. We investigated different 3D bioprinted GelMA constructs, i.e., a simple slab geometry with only NPs (Slab_NP), only algae (Slab_alg), as well as constructs with both algae and sensor NPs in a slab (Slab_NP+alg) or Vgroove geometry (Slab_NP+alg).

### 3.1 Growth measurements

We investigated the cell growth and photosynthetic capacity of the green microalgae in four identical bioprinted constructs (slab_NP+alg) (Figure 2). The printed constructs exhibited a relatively low maximum PSII quantum yield (F_v_/F_m_) of 0.48 on day 2 (Figure 2a), indicating that the microalgae were stressed from the bioprinting and curing process. Also the algal cell count did not increase and remained around 100 ± 43 x 10^6^ cells mL^-1^ (mean ± SD) during the first 2 days of the experiment (Figure 2b), probably due to effects of photocuring of the hydrogel matrix causing photo-bleaching of the microalgae in combination with slow growth in the low light during incubation. Right after curing, the outer layers of the bioprint progressively faded in color up until a few hours after printing due to loss of chlorophyll from damaged cells (Figure S2), while the cells recovered and exhibited faster growth after 2 days. The algal cell count increased exponentially to 320 ± 24 x 10^6^ cell mL^-1^ on day 8, while the F_v_/F_m_ increased to 0.65 indicating good algal health and photosynthetic efficiency.^85^

**Figure 2:**
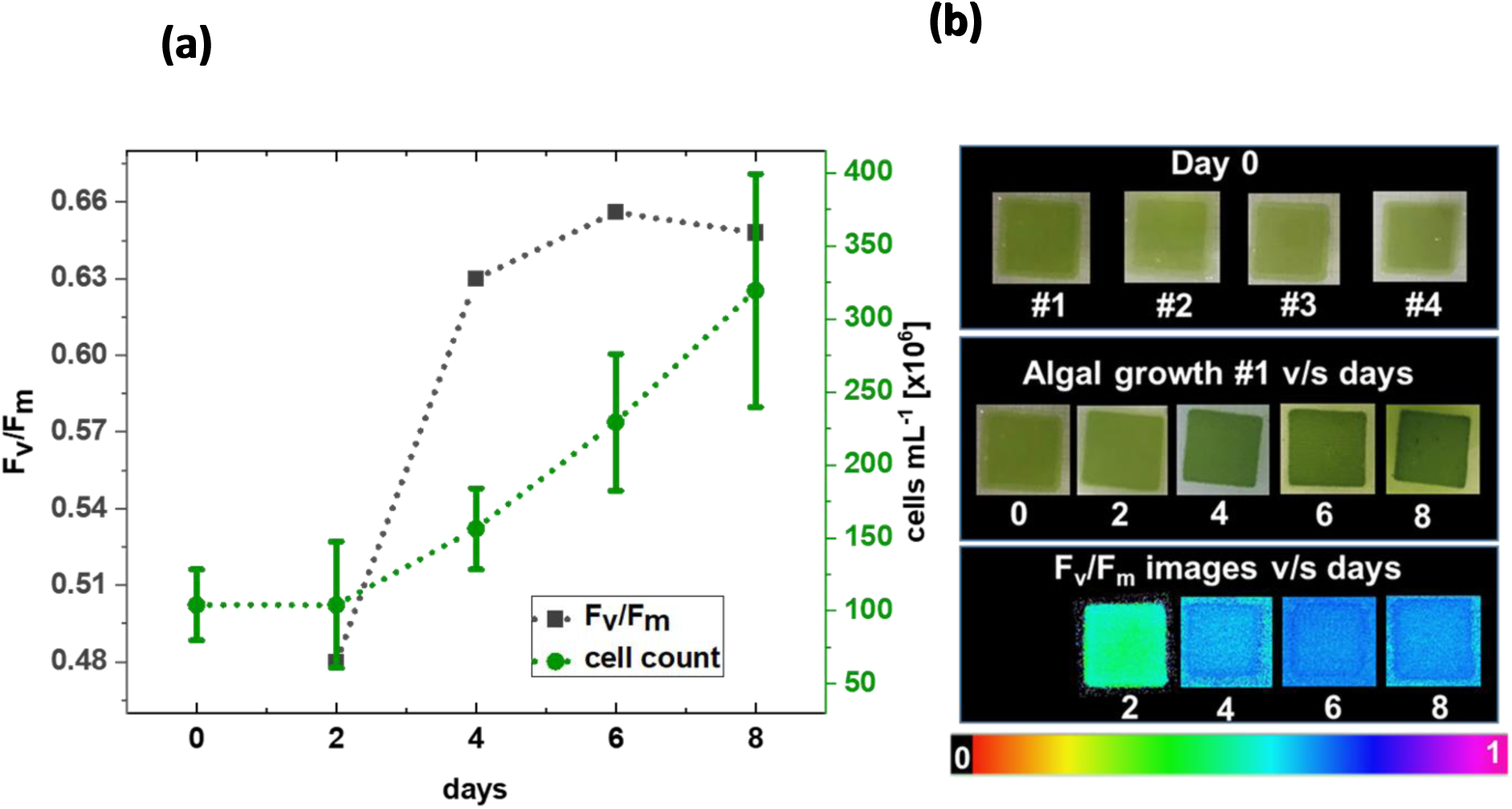
Cell growth and photosynthetic capacity in bioprinted constructs. (a): Cell counts and the maximum quantum yield of PSII activity (F_v_/F_m_) of green microalgae in 3D bioprinted GelMA constructs with optical O_2_ sensor NPs, from day 0 (cell counts were done on bioink just before printing) until 8 days after printing; (b): Photographs showing 4 identical bioprinted hydrogel slabs with microalgae and sensor NPs (Slab_NP+alg; #1 - 4, top row), algal growth in slab #1 over 8 days (middle row), and corresponding images of F_v_/F_m_ measured every 2 days during the growth phase (bottom row; false color coded according to the scale bar).

### 3.2 Photosynthetic performance of bioprinted constructs

We investigated the photosynthetic performance of 8-day-old algal bioprints by imaging the effective PSII quantum yield (YII) and the derived relative PSII electron transport rate (rETR) across the whole construct as a function of incident photon irradiance (Figure 3). We found that the addition of NPs and the topography of the bioprint (Vgroove versus slab) did not significantly affect the fitness of the microalgae, as all samples had similar F_v_/F_m_ values (∼0.65) and exhibited similar curves of YII and rETR versus photon irradiance (PAR; 400-700 nm) (Figure 3a). With increasing photon irradiance, the photosystems of the microalgae became increasingly saturated and consequently YII decreased. The rETR vs. photon irradiance curves indicate the onset of photosynthesis saturation at a photon irradiance of ∼250 μmol photons m^−2^ s^−1^. Images showed a relative uniform distribution of YII in bioprinted slabs with (Slab_NP+alg; Figure 3b top row) and without NPs (Slab_alg; Figure. 3b bottom row), the bioprinted construct with grooves (Vgroove_NP+algae; Figure 3b middle row) showed lower YII values in between the ridges of the construct, indicative of higher microalgal activity and growth on the ridges. The good biocompatibility of the functionalized GelMA bioink with sensor NPs is in line with an earlier study showing good photosynthetic performance of microalgae immobilized in a alginate/methyl-cellulose bioink with NPs.^31^

**Figure 3:**
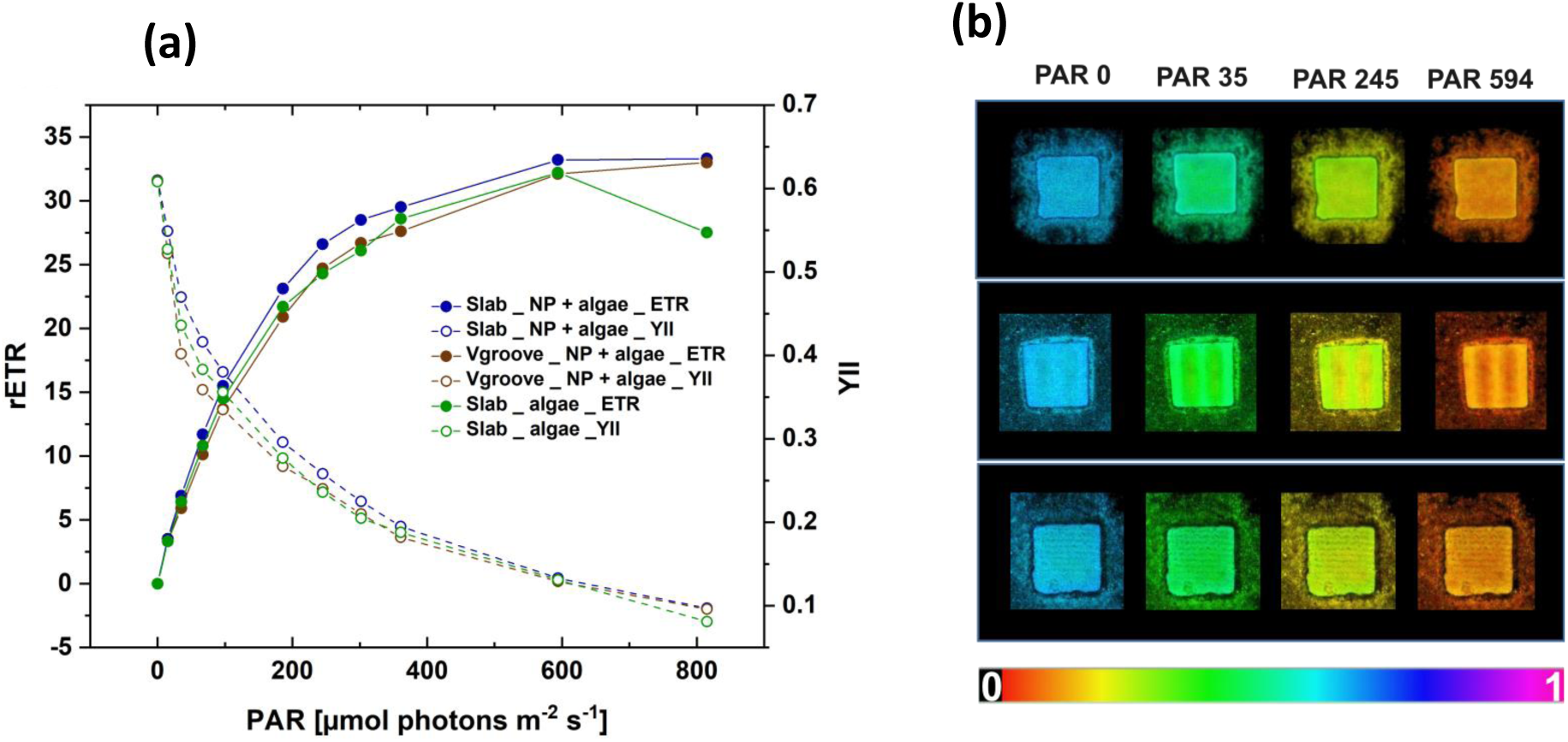
Photosynthetic performance of 8 day old algal bioprints: (a): Effective PSII quantum yield (YII) and the derived relative photosynthetic electron transport rate in PSII (rETR) versus photon irradiance (μmol photons m^−2^ s^−1^; PAR, 400-700 nm), as integrated over the entire sample; (b): Images of YSII measured at increasing incident photon irradiance levels of blue actinic light (460 nm). The values are false color coded according to the scale bar. Top: Slab_NP+algae; middle: Vgroove_NP+algae; bottom: Slab_algae.

### 3.3 Optical coherence tomography imaging of bioprinted construct

Optical coherence tomography (OCT) is an interferometric imaging technique using near infrared radiation for noninvasive 3D imaging of biological samples.^49^ It measures patterns of direct, elastically, backscattered photons, from surfaces with refractive index mismatches and from internal microstructures.^86^ It has been widely applied in dermatology and ophthalmology^49^ and in a few application for studying aquatic animal tissues with microalgal symbionts^87,88^, microalgal biofilms^48,89^. OCT has also been applied to study bacterial growth in alginate beads^90^ and to image natural corals for designing bioprinted bionic corals^9^, and e.g. to assess and optimize the print fidelity of construct designs in extrusion bioprinting^91^. To our knowledge this study is the first application of OCT for studying optical properties of bioprinted constructs.

OCT imaging provides information about scattering properties of the imaged samples. By fitting the depth dependent OCT signal attenuation and local OCT signal intensity, based on a theoretical model developed from inverse Monte Carlo modelling of photon transport, the scattering co-efficient (µ_s_) and anisotropy of scattering (g) can be extracted.^70^ In this study we apply this technique to image the microalgal distribution in 8-day-old bio-prints and extract the optical properties to be used for light simulations of radiative transfer in the bioprinted constructs (see section 4.7) (Figure 4). Figure 4 a-d, show the plot of focus-corrected, calibrated reflectivity values versus the sample depth, as obtained from cross-sectional OCT scans (OCT-b scans) shown as inserts. The exponential fit parameters (µ and *ρ*) for the 4 different samples were mapped to the scattering coefficient, µ_s_, and the scattering anisotropy, g, at 930 nm (Figure 4e). Figure 4f, shows the 3D rendering of the OCT image stacks for Slab_NP+alg and Vgroove_NP+alg. Signal attenuation along the depth of the samples can be observed.

**Figure 4:**
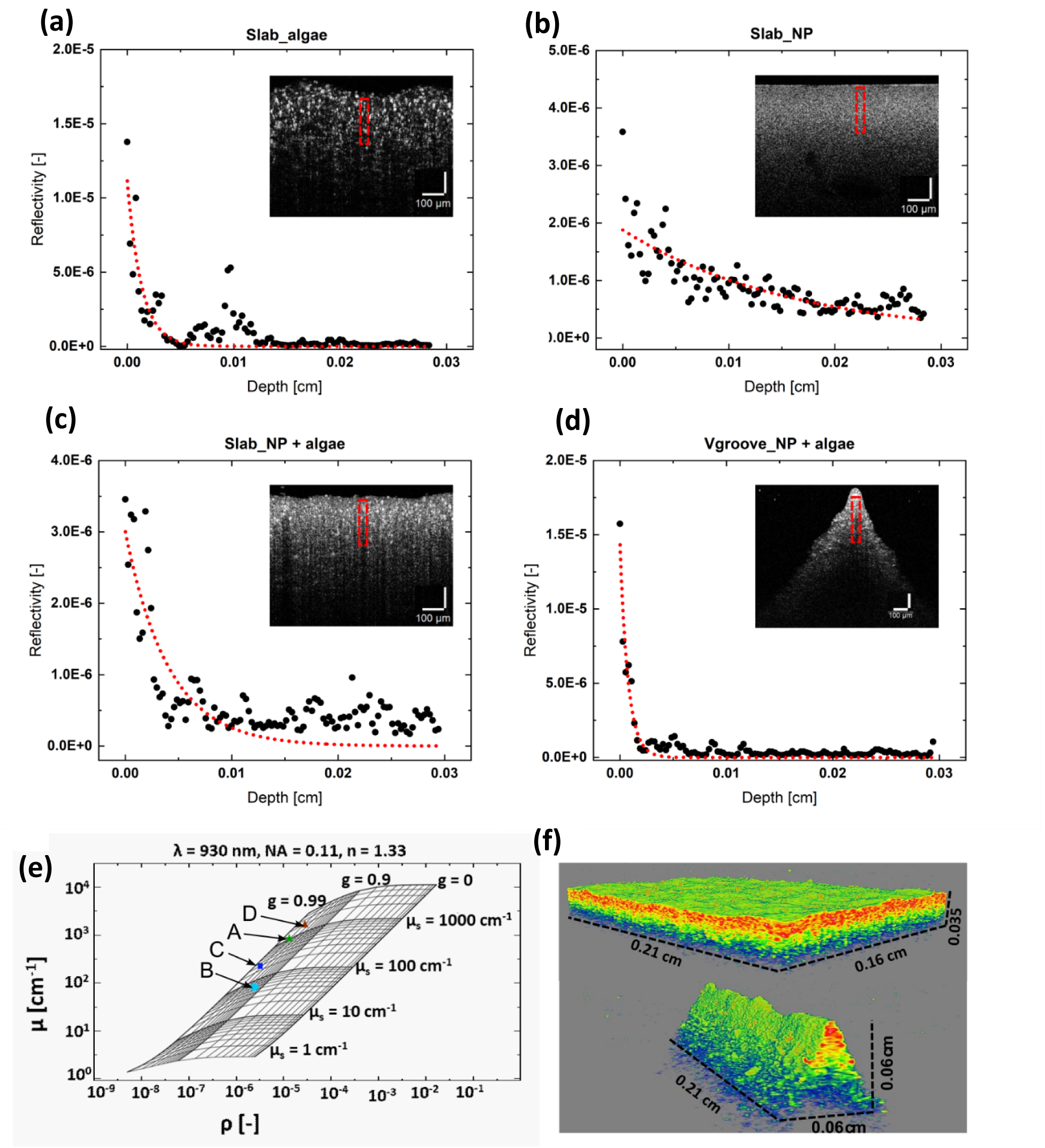
Optical coherence tomography imaging of 8 day old algal bioprints: a – d: Calibrated OCT signal fall-off with depth (averaged over a few pixels, along the horizontal direction, as indicated by the red box in the inserts showing corresponding cross-sectional OCT images) and the corresponding exponential fit for the different bioprint samples (a. Slab_alg, b. Slab_NP, c. Slab_NP+alg, d. Vgroove_NP+alg). E: Interpolation grid mapping of the experimental fit parameters (from a – d) to the corresponding material optical properties, i. e., µ_s_ and g at 930 nm. f: 3D rendering of the OCT image stacks of a Slab_NP+alg and Vgroove_NP+alg bioprinted construct, corresponding to samples shown in (c) and (d). Scale bar: 100 µm. The false color coding represents the intensity of the non-calibrated OCT signal, which was manually adjusted to optimize visualization of the structural features for each scan.

The extracted optical properties were used in the light simulation (section 2.4), solving the radiative transfer equation based on a Monte Carlo modeling approach^55,76^ for light at 636 nm, which is in the absorption range of Chl *b* in the green algae^92^. The optical parameters are summarized in Table S3. The absorption co-efficient (µ_a_) was obtained by cell counting, as described in section 4.11. The estimated µ_s_ from OCT measurements for Slab_alg was ∼1000 cm^-1^, whereas an estimation of µ_s_ based on sample cell count^78^ (described in methods section 4.7) was ∼285 ± 77.2 cm^-1^. This difference between the 2 methods can be explained by the fact that the cell count method^78^ estimates optical properties of cells suspensions and does not take algal clustering into account, which can significantly alter the scattering properties of the sample.

All 3 bioprinted constructs with algae had a g value of 0.98 indicating strong forward scattering. In contrast, Slab_NP had a g value of 0.93 indicating that the scattering from NP in GelMA is slightly more diffuse than from algal clusters. The Slab_alg had a higher µ_s_ compared to Slab_ NP+alg, 1000 cm^-1^ and 300 cm^-1^, respectively. The Slab_alg showed a steeper signal drop with depth in the construct (Figure 4 a) as compared to Slab_ NP+alg (Figure 4c). This is most likely because the NPs rendered the internal light field more diffuse, as seen in Slab_NP (Figure 4b). Addition of the NPs thus resulted in a more uniform internal light field. The OCT signal analysis for the Vgroove_NP+alg (Figure 4d) was conducted around the peak. Though the cell count for Vgroove_NP+alg and Slab_NP+alg is very similar, µ_s_ is 2000 cm^-1^ and 300 cm^-1^, respectively. The signal fall-off for the Vgroove_NP+alg was also steeper compared to Slab_NP+alg. We assign this difference to a higher algal density on the ridges of the construct, as compared to the rest of the Vgroove_NP+alg and the Slab_NP+alg constructs.

### 3.4. Mapping of O_2_-dynamics in bioprinted constructs

Luminescence lifetime imaging and respirometry measurements were conducted simultaneously, to quantify changes in O_2_ concentration due to microalgal photosynthesis and respiration in the bioprinted constructs. This combined approach provides i) 2D information of the O_2_ dynamics at the print-water interface (lifetime imaging, Figure 5), and ii) a measure of the total net O_2_ exchange across the whole construct surface (Figure 6 a-b) under defined levels of incident photon irradiance. The latter information was used to determine the total net production, dark respiration and gross photosynthesis (Figure 6 a-d) per area of the bioprinted constructs. Whereas the dissolved O_2_ concentration in the bulk water changes relatively slow depending on i) the water volume relative to the surface area of the bioprinted construct and ii) the flow-dependent mass transfer between construct and surrounding water, lifetime-imaging captured near real-time changes in dissolved O_2_ concentrations directly at the surface of the construct and revealed spatio-temporal heterogeneities in O_2_ dynamics (Figure 5, S5).

**Figure 5:**
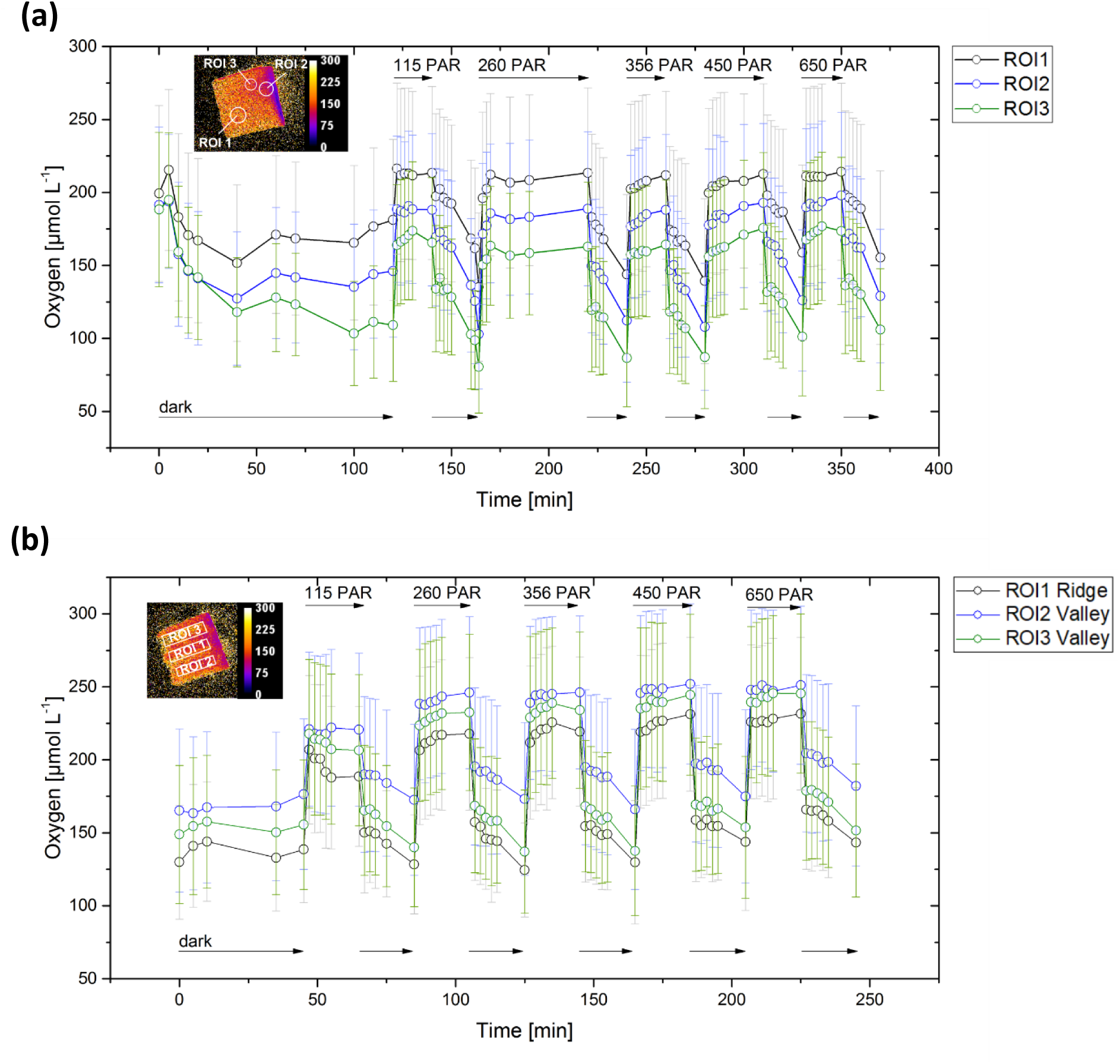
Oxygen dynamics at the bioprinted construct surface in 3 ROIs (shown in the inserts) for different light-dark cycles in the slab (a) and the v-groove (b) design. Three random ROIs were chosen for the slab, where as the v-groove ROI 1 was chosen along a ridge of the construct and ROI 2 and 3 in the surrounding valleys. Symbols with error bars represent means ± standard deviation within each ROI.

**Figure 6:**
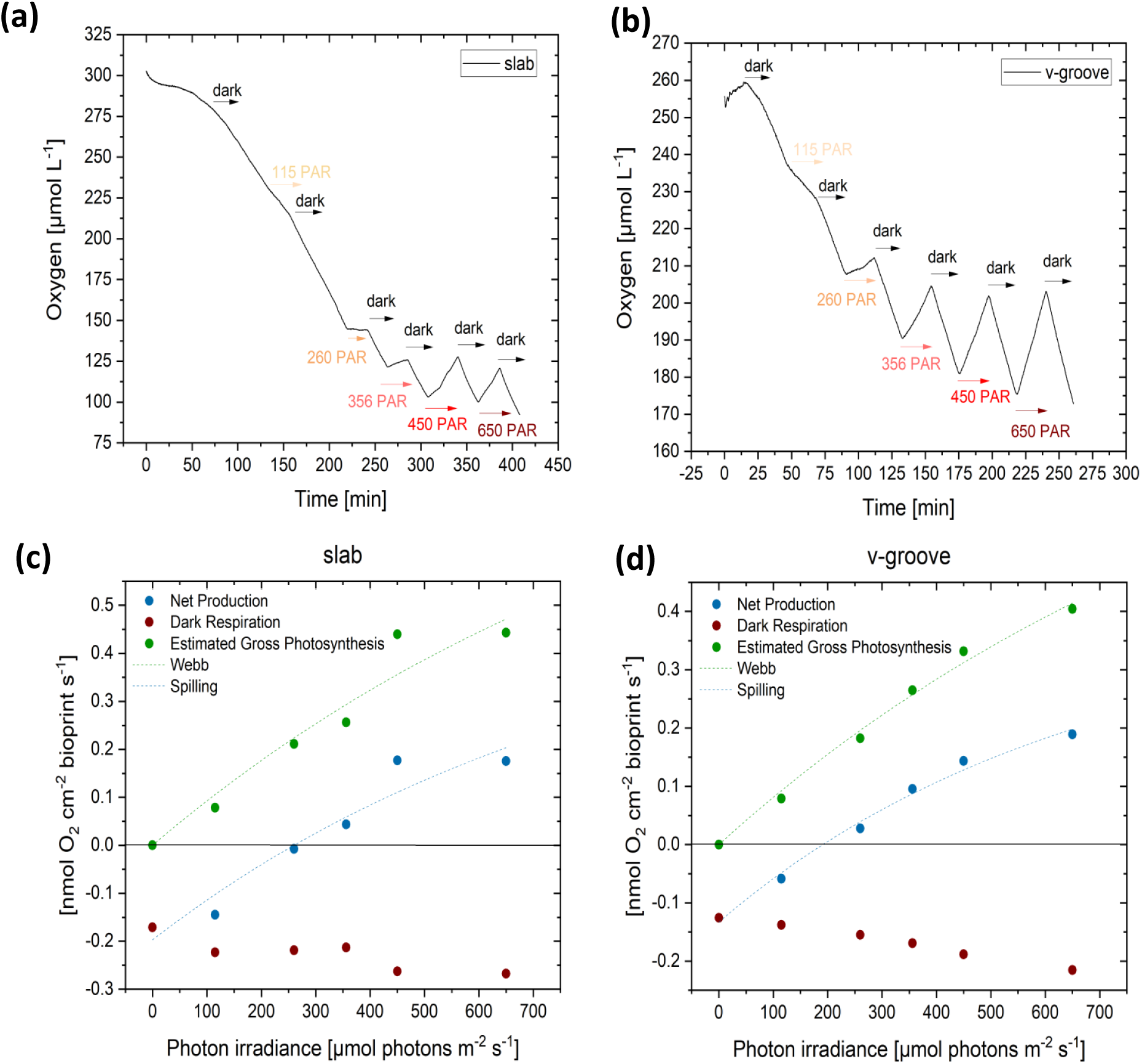
Gas exchange measurements of total photosynthesis and respiration of bioprinted constructs. Changes in O_2_ concentration measured over time under darkness and different light conditions with the slab (a) and the v-groove (b) bioprinted construct inside a gas-tight chamber. The rate of O_2_ change was determined from the linear slopes in (a) and (b), and were used to determine the net production, dark respiration and gross photosynthesis shown in (c) and (d).

In order to follow O_2_ dynamics at the surface of the print, the GelMA bioink was functionalized with O_2_-sensitive sensor nanoparticles. Changes in the dissolved O_2_ concentration cause a change in the luminescence lifetime of the sensor particles, which was imaged with a time-domain luminescence lifetime camera system (PCO SensiCam-SensiMod^53^), which is not affected by background luminescence (such as Chl fluorescence from the algae) or by uneven particle distribution in the sample.^54,82^ Calibration curves (Figure S6) showed no impact on the luminescence signal caused by GelMA, due to the optical clarity of the material. Lifetime images were acquired every 2 minutes for 10 minutes with an additional measurement after 20 minutes (Figure 5), whereas O_2_ in the respirometry chamber was logged continuously every 6 seconds (Figure 6 a-b). Theoretically, lifetime images can be acquired even more frequently^53^, however, measurements have to be done in darkness, thus the light needs to be briefly switched off for each measurement. Therefore, two-minute intervals were chosen as compromise between the amount of acquired images and the time the light had to be switched on and off – which can influences O_2_ production and consumption as well.

Lifetime imaging showed a rapid change of the O_2_ concentration at the bioprint – water interface reaching stable values well within the first 2 minutes of illumination (potentially much faster), with maximum changes of ∼70 µmol L^-1^ (slab) and 97 µmol L^-1^ (v-groove) in light within the first two minutes and drops of up to 43 µmol L^-1^ (slab) and 76 µmol L^-1^ (v-groove) in darkness. Overall, the v-groove design showed stronger dynamics in all ROIs in comparison to the slab, which can be attributed to the increased surface area enhancing both mass transfer and light harvesting. Respirometry showed similar, but slower dynamic changes in O_2_ concentration in the water surrounding the construct (Figure 6).

Lifetime imaging showed increases in dissolved O_2_ concentration for both geometries at the print-water interface already at the lowest used irradiance (115 PAR, Figure 5); whereas, the total O_2_ flux measurements showed net production in the v-groove at light intensities from ≥ 260 PAR (Figure 6b) and ≥ 356 PAR for the slab geometry (Fig. 6A). This could be due to better light distribution in the v-groove than the slab, or the increased surface area of the v-groove in comparison to the slab. Overall, the v-groove designed showed a more dynamic response to changes in the light conditions than the slab design, emphasizing the importance of surface topography for optimizing productivity in 3D constructs.

#### Light and mass transfer simulations

A simulation of light availability in the algal bioprints was be used in combination with diffusion-reaction models and laminar flow simulation to determine net O_2_ exchange and the steady state distribution of O_2_ concentration within the printed constructs under defined light conditions. This approach was first developed for multiphysics modeling of the radiate, heat and mass transfer in a multilayered coral model.^51^ Calculating both the light distribution and mass transfer can aid in predictive modeling to design efficient topographies for enhanced algal growth and help gain deeper understanding of the experimental data and observations.

First we calculated the light field in the bioprints using a finite element Monte Carlo (MC) modelling approach using unstructured meshes of tetrahedral elements, which enables discretization of complex geometries and result in a better accuracy of the photon transport solution than 3D voxel discretization.^93,94^ The results of the light simulation on the algal bioprints on day 0 and day 8 are shown in Figures 7a and d respectively. The assumed optical properties are summarized in Table S1. For day 0, the optical properties were calculated based on algal cell counts in the print, whereas for day 8 they were obtained from OCT images to account for the algal cell clustering during growth (see section 2.7 and 2.10).

**Figure 7:**
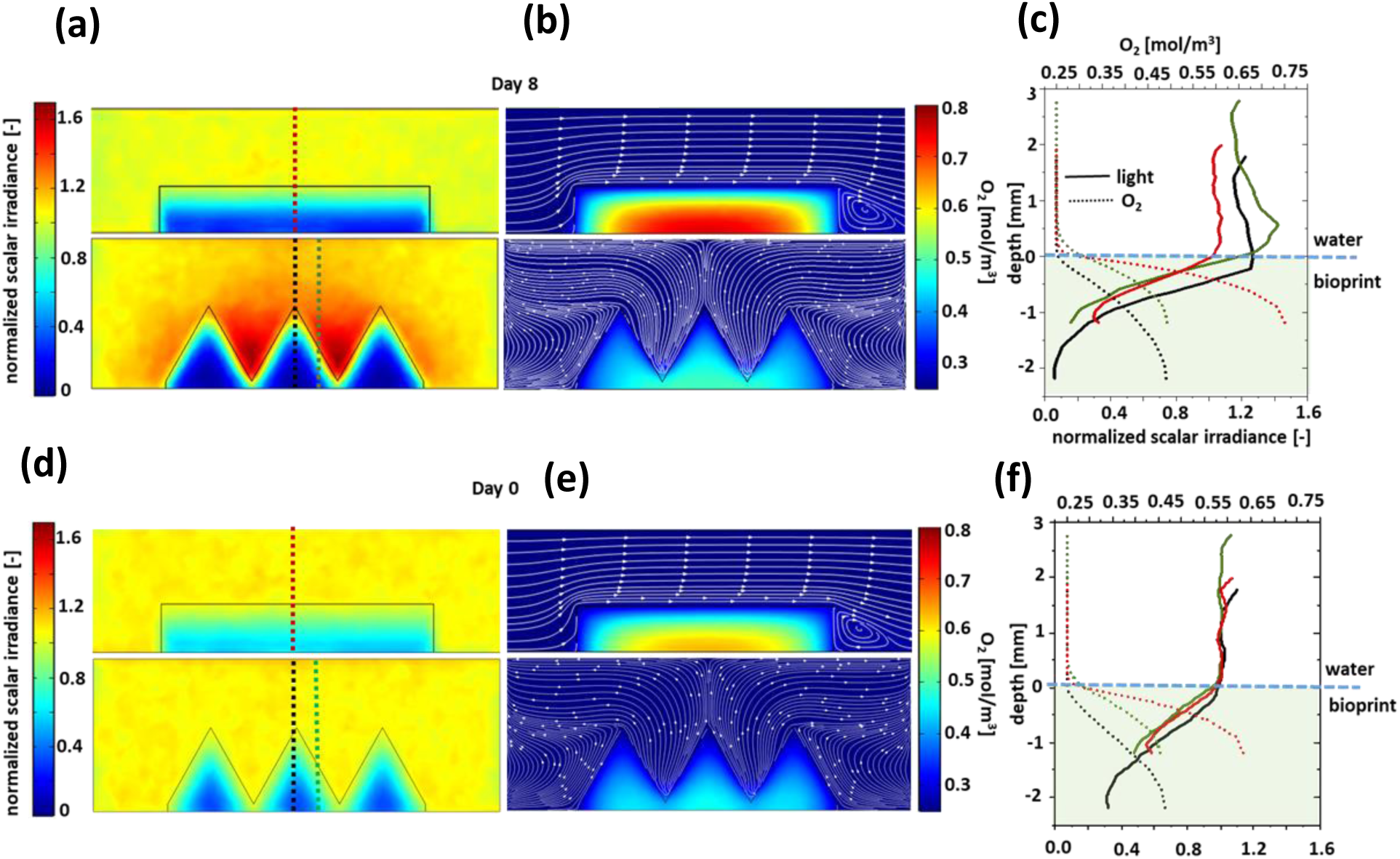
Simulated scalar irradiance, flow pattern and oxygen distribution in bioprinted constructs. (a) and (d): 2D light fields on day 8 and day 0, respectively, extracted from the central plane in the 3D model (see Figure S16). (b) and (e): 2D oxygen concentration distribution on day 8 and day 0, respectively, extracted from central plane in the 3D model (Figure S16). The white lines show the laminar flow pattern of water around the bioprints. (c) and (f): 1D line profiles of light and oxygen concentration extracted along the dotted lines as shown in A and D (red: center slab, black: center v-groove, green: slant v-groove). Note the different color scales for Day 0 and Day8, which were chosen to visualize more details in the data sets.

The light simulations indicate a higher light availability along the ridges of the v-groove construct than in the flat slab model, especially on day 8 (Figure 7). However, along the slopes of the ridges in the v-groove, the light intensity is reduced due to increased surface reflections under increasing incidence angle of light. Since the algal biomass is lower on day 0, more light reaches deeper into the bioprinted constructs on day 0 than on day 8 (Figure 7a, d). In spite of an increased illuminated surface area in the v-groove construct, the total photon flux density for a given incident irradiance in the v-groove was about 9% less than that in the slab (Table S4) on both day 0 and day 8, which can again be due to loss of light at the slant edges of the v-groove.

The simulated scalar irradiance line profiles along the centre of the slab, especially on day 8 (Figure 7C), is comparable to light micro profile measurements in natural algal and cyanobacterial biofilms^77^, where light typically attenuates strongly within the upper 1-2 mm of the biofilm, due to strong scattering and absorption by pigmented photosynthetic microbes. The absence of any distinct peaks in the light profile and penetration of light up to 1 (to 2) mm in the prints (except for v-groove on day 8) can be attributed to the forward scattering properties of the bioprint (g = 0.98). However, for the v-groove sample on day 8 a distinct peak in the scalar irradiance line profile can be observed, especially along the slopes (Figure 7c), which is due to increased back scattering with increasing angle of incidence of the incoming light.

The mass transfer simulations on both day 0 and day 8 (Figure 7b and e, cut line profile in Figure 7 c and f, respectively) show that there is less O_2_ accumulation in the v-groove design, as compared to the slab. The higher surface area to volume ratio for the v-groove (∼ 1.42 times the S/A ratio of the slab), results in a more efficient mass transfer with the surrounding medium, and the v-groove has 18.6 % and 33.6% less O_2_ in the bioprinted construct on day 0 and 8, respectively, as compared to the slab (Table S4). The enhanced O_2_ mass transfer in the v-groove construct on day 8 (versus day 0) is due to lesser O_2_ production deeper in the bioprint. The relatively higher biomass on day 8 leads to decreased light penetration deeper into the v-groove. As a consequence O_2_ production is limited to the region closer to the surface of the bioprint. This enables efficient diffusion of excess O_2_ from the vgroove to the surrounding medium. The O_2_ mass transfer simulation can be an indirect indicator for the mass transfer of other species like CO_2_ and nutrients, which are essential for the algal growth. 2D cross-section plots of light field and O_2_ concentration for the v-groove, along the length of the flow chamber are shown in Figure S11. However, direct comparison between the simulated O_2_ profiles in the bioprints and those measured in natural biofilms can be challenging because of the high variability in morphology and constitution of natural biofilms.

The surface plots of the simulated O_2_ concentration for slab and v-groove (Figure S14) and the O_2_ measurements of the samples shown in Figure 5 are not very comparable. One of the main reasons is that the mass transfer simulations were performed assuming a laminar flow around the samples, whereas the O_2_ measurements were conducted in closed vials with stirring at the bottom of the sample holder. Consequently, medium flow around the bioprinted constructs might have been different in the measurements e.g. due to some turbulence, as compared to the simulated laminar flow velocity in the multiphysics model. In order to make direct comparisons between simulations and measurements, future experiments should thus be designed with similar boundary and system conditions as in simulations and vice-versa.

## 4 Conclusion and Outlook

3D bioprinting provides a rapid prototyping platform to study complex phenomena involving the interaction of light and matter in biological systems and living cells. It can be combined with other high resolution fabrication techniques like two-photon polymerization^95^ and lithography^9^ to replicate complex topographies and hierarchical structures observed in nature facilitating incorporation of new sustainable biomimetic functionalities to products and devices.In this study we present a fabrication, imaging and simulation pipeline to evaluate photo-physiological performance of 3D bioprinted microalgal constructs, which have a direct application in designing optimal topographies enhancing algal growth, e.g. for biofuel and secondary algal metabolite production. This overcomes the limitation of currently used monitoring techniques, where cell metabolism and performance is measured in the media surrounding the constructs or by destructive sample analyses. Hence, this approach allows for in-line, non-invasive motoring of the bioprinted samples and provides effective strategies for design optimization based on a combination of mechanistic model predictions and experimental validation to achieve high photosynthetic efficiencies and growth rates. The presented pipeline can also be extended to other areas of life sciences facilitating rapid evaluation of cell activity as a function of structural complexity in printed constructs, metabolic interactions in mixed species bioprints, and in response to external incubation conditions.

The combination of 3D bioprinting with non-invasive functional imaging (e.g. based on inclusion of optical sensor particles in constructs), respirometry and multi-physics simulation provides a useful toolkit to simultaneously study gradients of light and O_2_ e.g. within synthetic biofilms, tissue constructs and artificial co-culture systems, while at the same time quantifying overall performance of bioprinted constructs with different geometries in terms of O_2_ production or uptake. Such a combined modelling and experimental validation approach could e.g. be used to monitor and optimize O_2_ supply to cells in 3D *in vitro* cell and tissue cultures, in order to achieve oxygenation of the entire culture.^42^ While we have demonstrated the use of functionalized bioink for non-invasive monitoring of O_2_ dynamics in bioprinted constructs via ratiometric^31^ and luminescence lifetime imaging (present study), bioinks can be functionalized with other types of optical sensor particles^96 62^, expanded to other parameters e.g. via optical sensing of temperature, pH and other chemical species like CO_2_ and NH_3_, which can provide further insights into cellular interactions, particular metabolic functions and growth constraints. Non-invasive 3D imaging technologies like light sheet microscopy^46^ and confocal imaging^42^ can in principle also be implemented to map biomass and O_2_ distributions simultaneously, to study structure-function relationships in bio-printed constructs at even higher spatial resolution. Future studies will involve fabrication and investigation of heterogeneous, stratified (and spatially separated) bionic structures, mimicking natural systems like corals ^9^, beach rock^97^ etc. This will aid investigation of microbial and host-symbiont interactions under relatively well defined conditions. It can also, to some extent, simplify the development of the imaging/sensing systems, by reducing the variability observed in natural samples.

## Acknowledgements

We thank Sofie Lindegaard Jakobsen and Veronica Bach Petersen for excellent technical assistance. The study was supported by research grants from the Villum Foundation [VIL50371 & VIL57413, MK], the Independent Research Fund Denmark [DFF-8022-00301B; MK], a grant from the European Union’s Horizon 2020 research and innovation program [Marie Skłodowska-Curie grant agreement No 860125], and the Gordon and Betty Moore Foundation [grant no. GBMF9206; https://doi.org/10.37807/GBMF9206; MK].

## Author Contributions

SM, MM and MK conceptualized and designed the research. SM and MM synthesized GelMA. SM prepared the bionk, optimized printing and photocuring, and fabricated the bioprints. SM performed OCT imaging and analysis, as well as cell counts in bioprinted constructs. MM performed combined lifetime and respirometry measurements and image analysis. SM performed numerical simulations. SM, MM and MK wrote the manuscript with editorial help from ET.

## Supplementary information

**Table S1:**
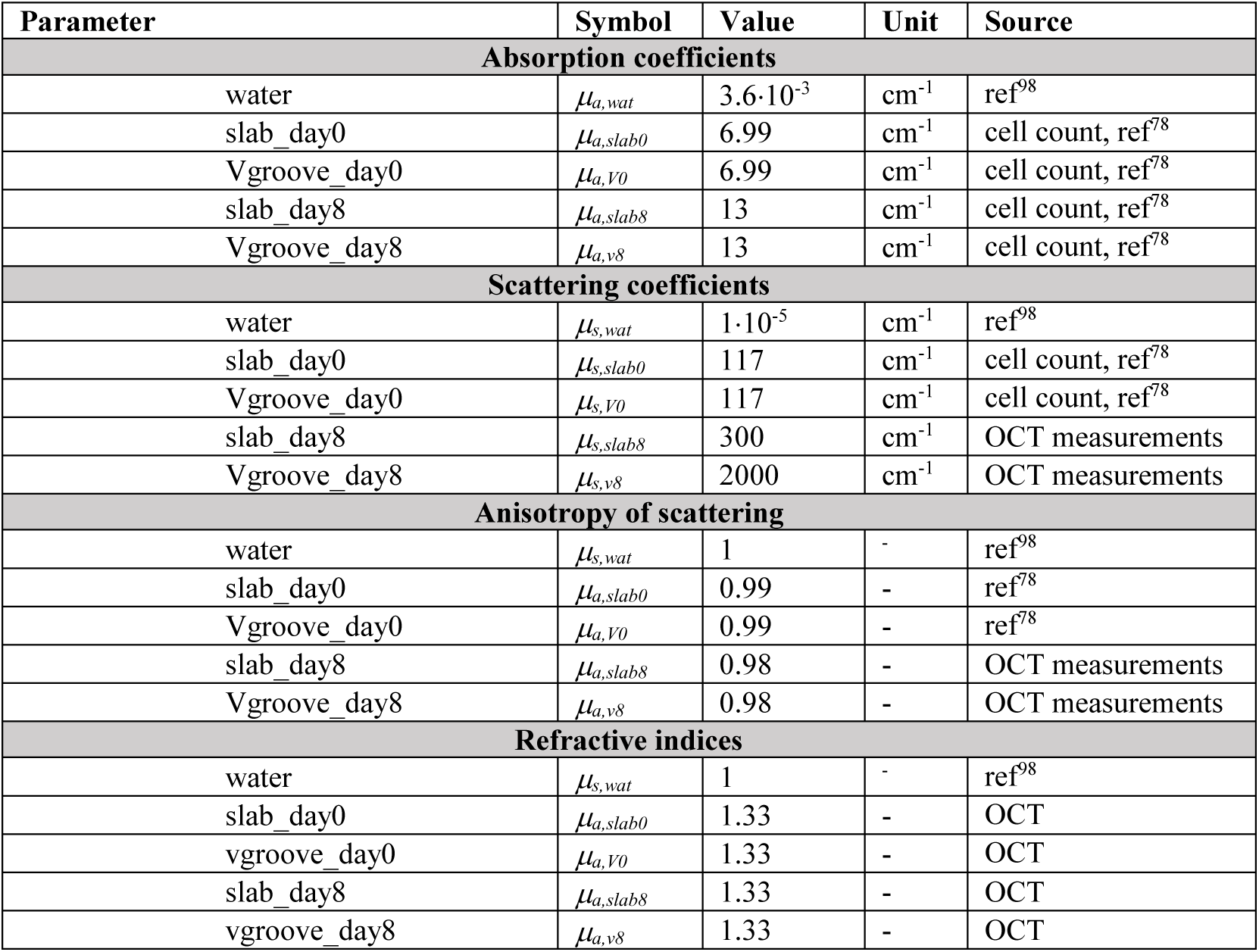
Model parameters used for MC simulations of radiative transfer (636 nm) in bioprinted slabs.

| Parameter | Symbol | Value | Unit | Source |
| --- | --- | --- | --- | --- |
| **Absorption coefficients** |  |  |  |  |
| water | $\mu_{a,wat}$ | $3.6 \cdot 10^{-3}$ | $\text{cm}^{-1}$ | ref<sup>98</sup> |
| slab_day0 | $\mu_{a,slab0}$ | 6.99 | $\text{cm}^{-1}$ | cell count, ref<sup>78</sup> |
| Vgroove_day0 | $\mu_{a,v0}$ | 6.99 | $\text{cm}^{-1}$ | cell count, ref<sup>78</sup> |
| slab_day8 | $\mu_{a,slab8}$ | 13 | $\text{cm}^{-1}$ | cell count, ref<sup>78</sup> |
| Vgroove_day8 | $\mu_{a,v8}$ | 13 | $\text{cm}^{-1}$ | cell count, ref<sup>78</sup> |
| **Scattering coefficients** |  |  |  |  |
| water | $\mu_{s,wat}$ | $1 \cdot 10^{-5}$ | $\text{cm}^{-1}$ | ref<sup>98</sup> |
| slab_day0 | $\mu_{s,slab0}$ | 117 | $\text{cm}^{-1}$ | cell count, ref<sup>78</sup> |
| Vgroove_day0 | $\mu_{s,v0}$ | 117 | $\text{cm}^{-1}$ | cell count, ref<sup>78</sup> |
| slab_day8 | $\mu_{s,slab8}$ | 300 | $\text{cm}^{-1}$ | OCT measurements |
| Vgroove_day8 | $\mu_{s,v8}$ | 2000 | $\text{cm}^{-1}$ | OCT measurements |
| **Anisotropy of scattering** |  |  |  |  |
| water | $\mu_{s,wat}$ | 1 | - | ref<sup>98</sup> |
| slab_day0 | $\mu_{a,slab0}$ | 0.99 | - | ref<sup>78</sup> |
| Vgroove_day0 | $\mu_{a,v0}$ | 0.99 | - | ref<sup>78</sup> |
| slab_day8 | $\mu_{a,slab8}$ | 0.98 | - | OCT measurements |
| Vgroove_day8 | $\mu_{a,v8}$ | 0.98 | - | OCT measurements |
| **Refractive indices** |  |  |  |  |
| water | $\mu_{s,wat}$ | 1 | - | ref<sup>98</sup> |
| slab_day0 | $\mu_{a,slab0}$ | 1.33 | - | OCT |
| vgroove_day0 | $\mu_{a,v0}$ | 1.33 | - | OCT |
| slab_day8 | $\mu_{a,slab8}$ | 1.33 | - | OCT |
| vgroove_day8 | $\mu_{a,v8}$ | 1.33 | - | OCT |

**Table S2:** Model parameters used in the simulations of flow, mass transfer and metabolism.

| Parameter | Symbol | Value | Unit | Source |
| --- | --- | --- | --- | --- |
| **Flow** |  |  |  |  |
| Viscosity water | $\eta_{wat}$ | 0.0009 | Pa s | ref<sup>99</sup> |
| **Densities** |  |  |  |  |
| water | $\rho_{wat}$ | 997 | kg m<sup>-3</sup> | ref<sup>99</sup> |
| bio-print | $\rho_{bp}$ | 1109 | kg m<sup>-3</sup> | ref<sup>100</sup> |
| **Mass transport** |  |  |  |  |
| **Diffusion coefficients** |  |  |  |  |
| O<sub>2</sub> in water | $D_{O_2, wat}$ | 2·10<sup>-9</sup> | m<sup>2</sup> s<sup>-1</sup> | ref<sup>101</sup> |
| O<sub>2</sub> in bio-print | $D_{O_2, bp}$ | 1·10<sup>-9</sup> | m<sup>2</sup> s<sup>-1</sup> | ref<sup>102</sup> |
| **Experimental parameters** |  |  |  |  |
| Average water velocity | $v_0$ | 5 | mm s<sup>-1</sup> | ref<sup>103</sup> |
| Ambient temperature | $T_0$ | 25 | °C | ref<sup>103</sup> |
| Inlet O<sub>2</sub> concentration | $c_{0, O_2}$ | 0.25 | mol m<sup>-3</sup> | ref<sup>104</sup> |
| Incident downwelling photon irradiance ( | $I_0$ | 430 | μmol photons m<sup>-2</sup> s<sup>-1</sup> | |

**Table S3:** Table summarizing the optical properties at 636 nm of bioprinted constructs extracted from OCT data on day 8 after incubation under a photon irradiance (400-700 nm) of 6 µmol photons m^-2^ s^-1^.

| Sample after 8 days | Number of cells (x 10<sup>6</sup>) | $\mu_s$ [cm<sup>-1</sup>] | $g$ | $\mu_a$ [cm<sup>-1</sup>] |
| --- | --- | --- | --- | --- |
| slab_NP | --- | 50 | 0.93 | --- |
| slab_alg | 14 ± 3.8 | 1000 | 0.98 | 17 |
| Slab_NP+alg | 11 ± 3.6 | 300 | 0.98 | 13 |
| Vgroove_NP+alg | 11 ± 4.1 | 2000 | 0.98 | 13 |

Table S4. Comparison of simulated light field (636 nm) and O_2_ content for bioprinted constructs designs with slab and Vgroove geometry, integrated over the entire volume.

**Figure S1:**
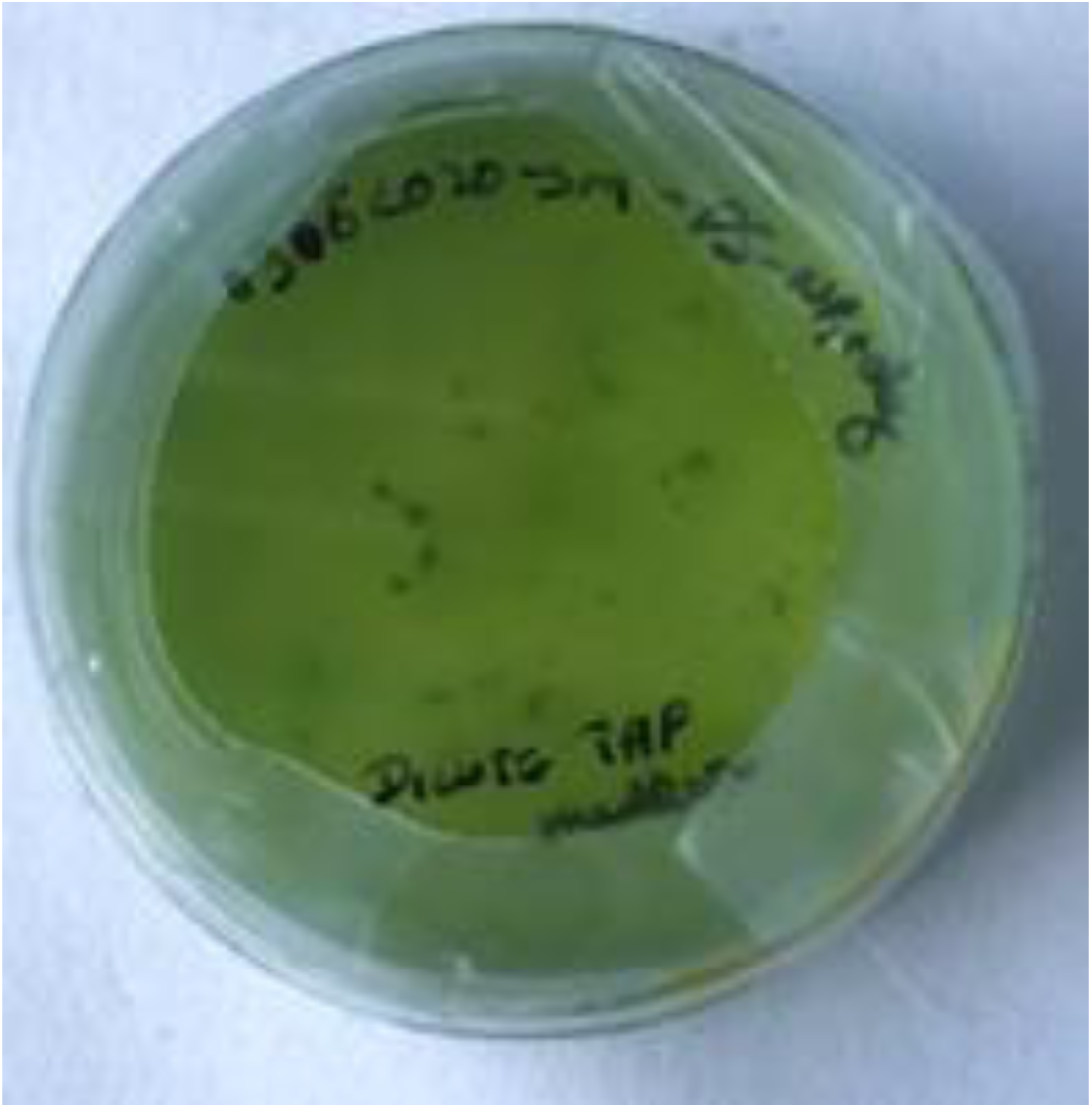
Photograph of degraded algal bioprint releasing the algae into the surrounding TAP medium.

**Figure S2:**
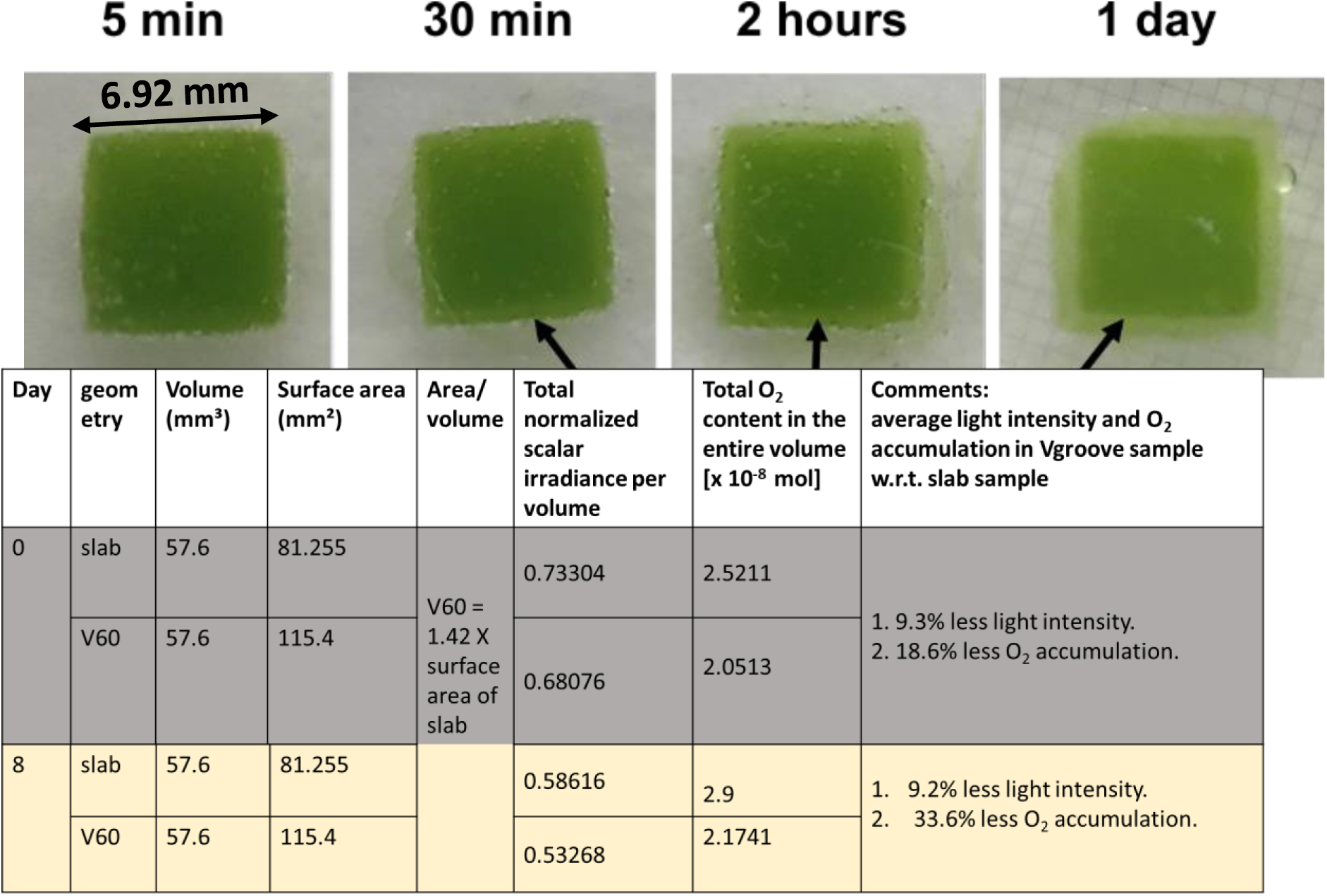
Photographs of an UV-cured (365 nm), 2.1 mm thick bio-printed construct with a simple slab geometry (Slab_alg) imaged in TAP medium at different times after curing. The slab was stored under an incident photon irradiance (400-700 nm) of 6 µmol photons m^-2^ s^-1^ over 1 day. The high UV dose required for curing the bioink resulted in bleaching and growth inhibition of microalgae in the outer layers of the construct as seen as the development of a bleached edge of the slab over time.

**Figure S3:**
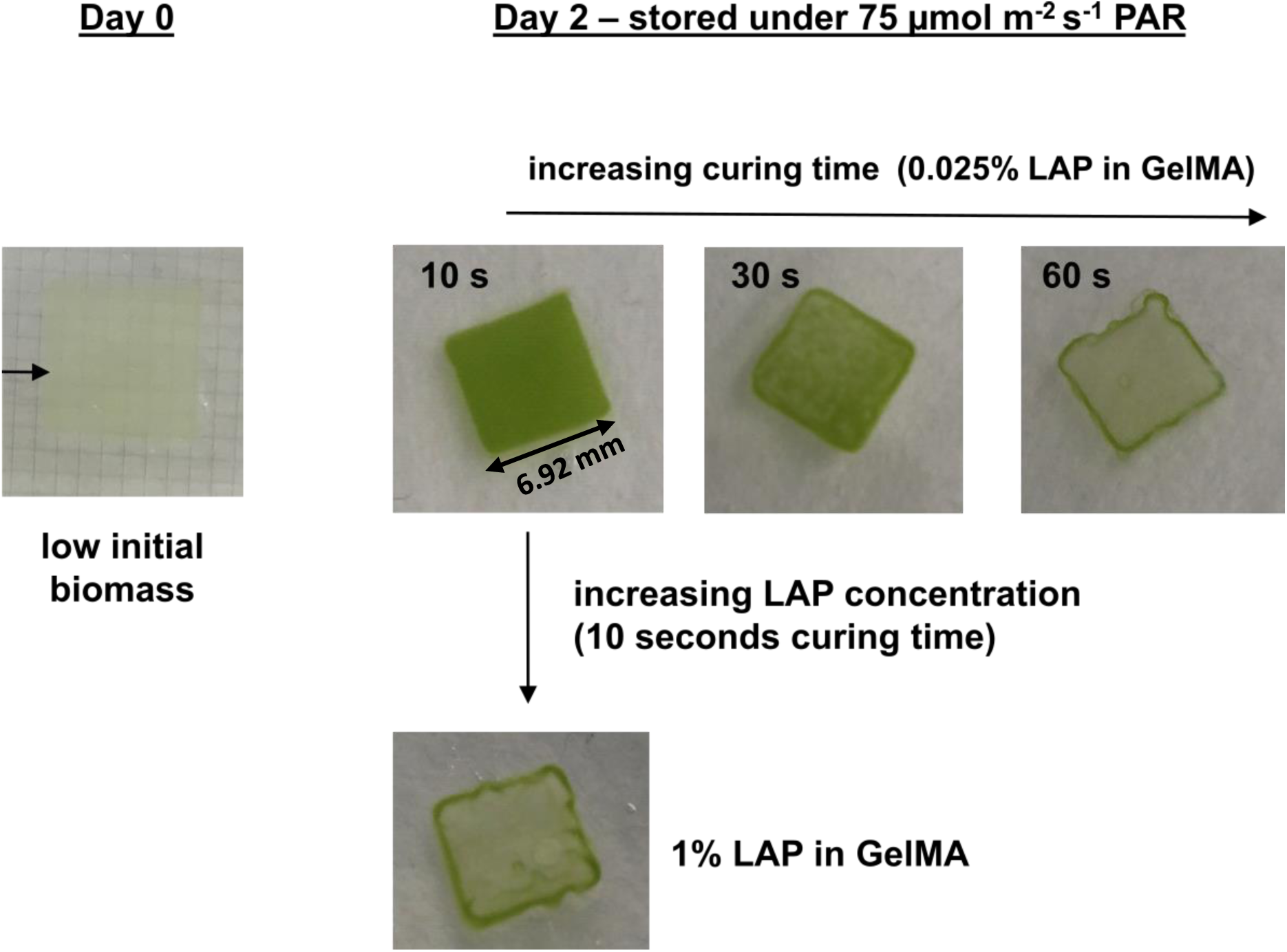
Photographs of a 1.1 mm thick bio-printed slab og GelMA with green microalgae and LAP photo curing agent for blue light curing (405 nm). Algal cell viability reduces with increasing curing dose and increasing LAP concentration. The optimized curing time for 405 nm light at 3 mWcm^-2^ was 30 seconds, using a LAP concentration (w/v) around 0.025%.

**Figure S4:**
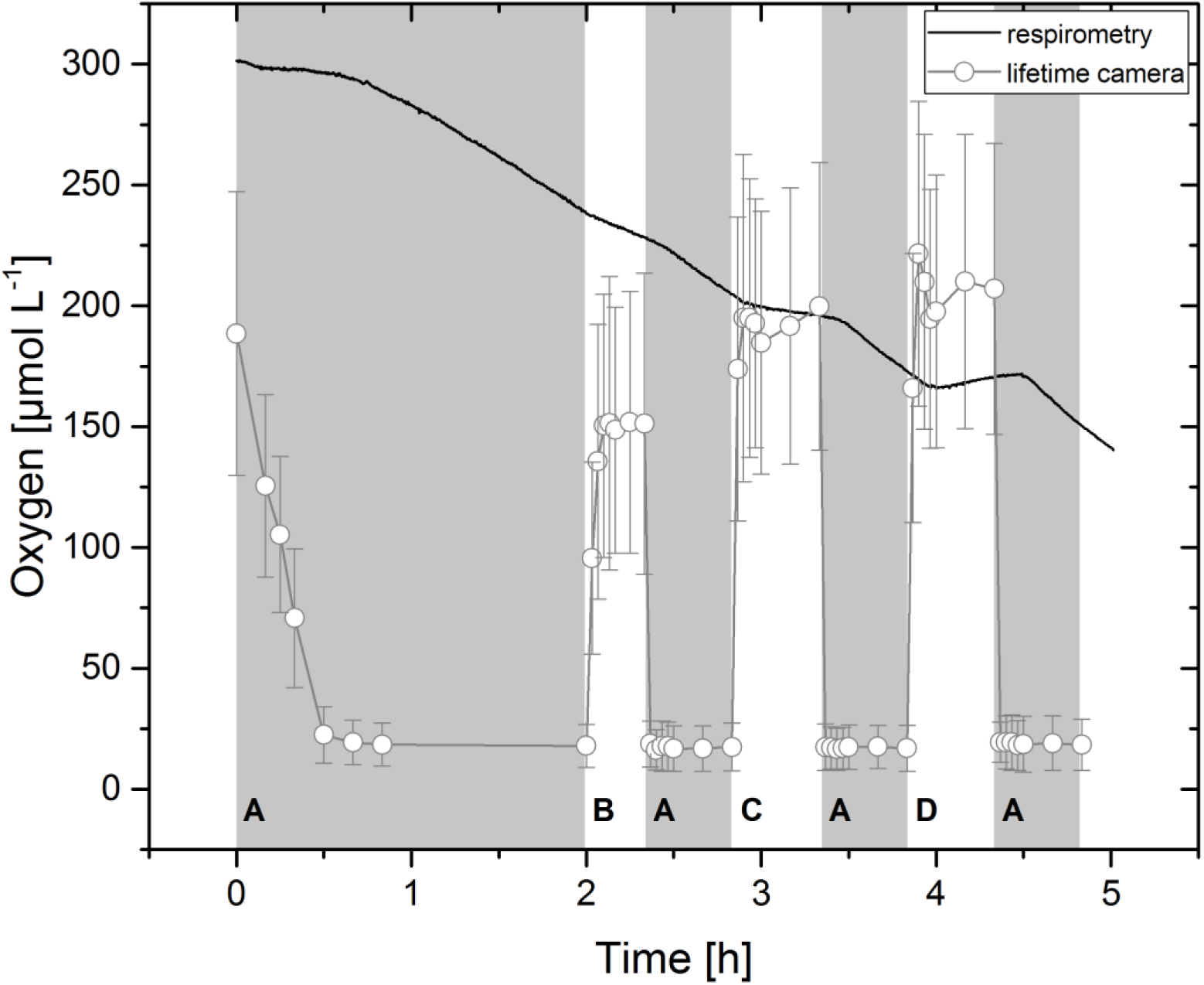
Simultaneous respirometry (black solid line) measurements of the net O_2_ exchange and luminescence lifetime imaging (grey line with symbols) of the surface O_2_ concentration on a bioprinted GelMA slab with microalgae and O_2_ sensor NPs. Respirometry showed a shift from net O_2_ uptake towards net O_2_ production at highest photon irradiance. Luminescence lifetime imaging could measure the fast O_2_ dynamics occurring at the slab surface. High error bars indicate uneven signal intensities, which can be attributed to local variations in O_2_ production and consumption. The grey background indicates darkness (A), while white areas represent measurements under different photon irradiance (400-700 nm), i.e., B = 80, C = 275, and D = 456 µmol photons m^-2^ s^-1^. Symbols±error bars indicate means±standrd deviation.

**Figure S5:**
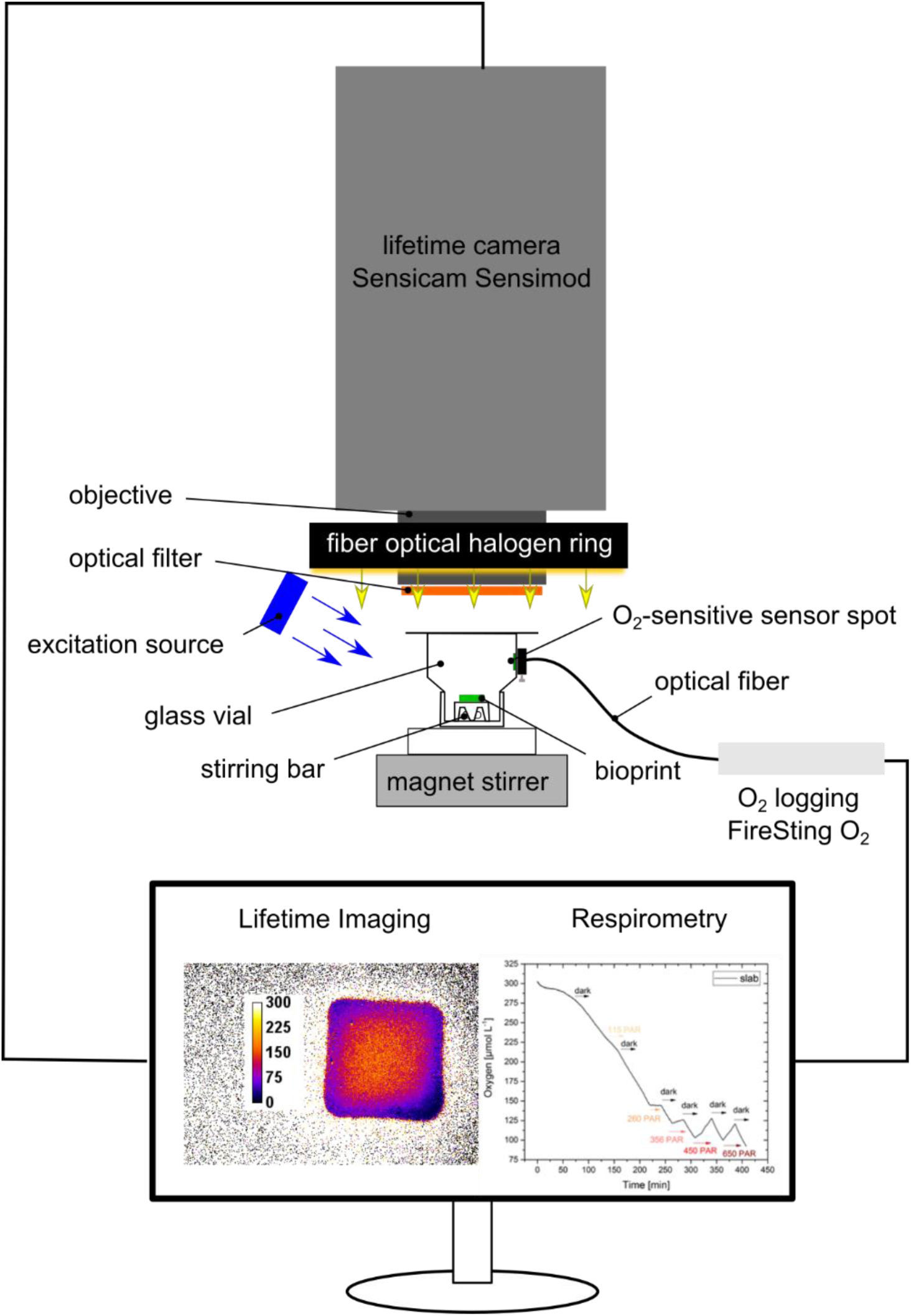
Experimental setup for combined respirometry and luminescence lifetime imaging of O_2_ dynamics. A bioprinted construct is placed inside a custom made, air-tight glass chamber on a small pedestal above a magnetic stirring bar, ensuring even mixing of the water in the chamber and efficient gas exchange between the bioprinted construct and the surrounding water. Light-dependent changes in total dissolved O_2_ within the chamber are measured with an O_2_-sensitive sensor spot that is read out with an optical fiber attached to the transparent chamber wall and connected to a fiber-optic O_2_ meter. Simultaneous 2D mapping of O_2_ concentration over the construct is done with a luminescence lifetime camera, placed perpendicular to the bioprint containing O_2_-sensitive sensor NPs. A 460 nm LED is used for excitation of the luminescent sensor particles. A long-pass filter (565 nm) is placed in front of the camera lens. Irradiance is controlled via a fiber optical halogen ring light, placed around the camera objective.

**Figure S6:**
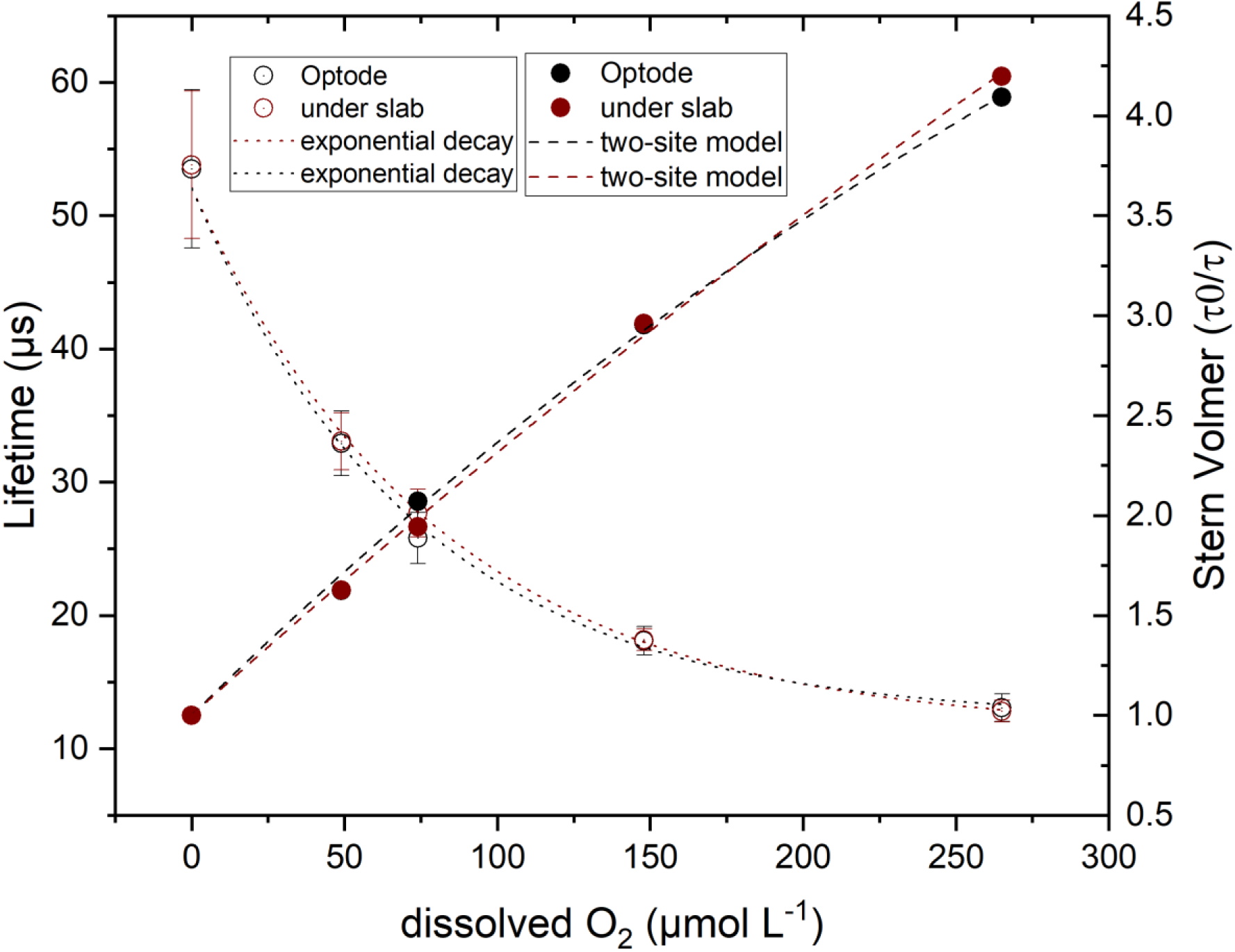
Calibration curves of the O_2_ sensitive indicator dye shown as the measured luminescence lifetimes (open symbols) and the normalized lifetimes (τ_0_/τ) (close symbols), as derived from imaging a planar optode (black symbols) or a clear bioprinted GelMA slab on top of the planar optode (red symbols). The luminescence lifetime vs. O_2_ concentration measurements were fitted with an exponential decay function (R^2^> 0.99) (dotted), while the normalized lifetime vs O_2_ concentration measurements were fitted with a modified Stern-Volmer equation using the two-site model^81,105^.

**Figure S7:**
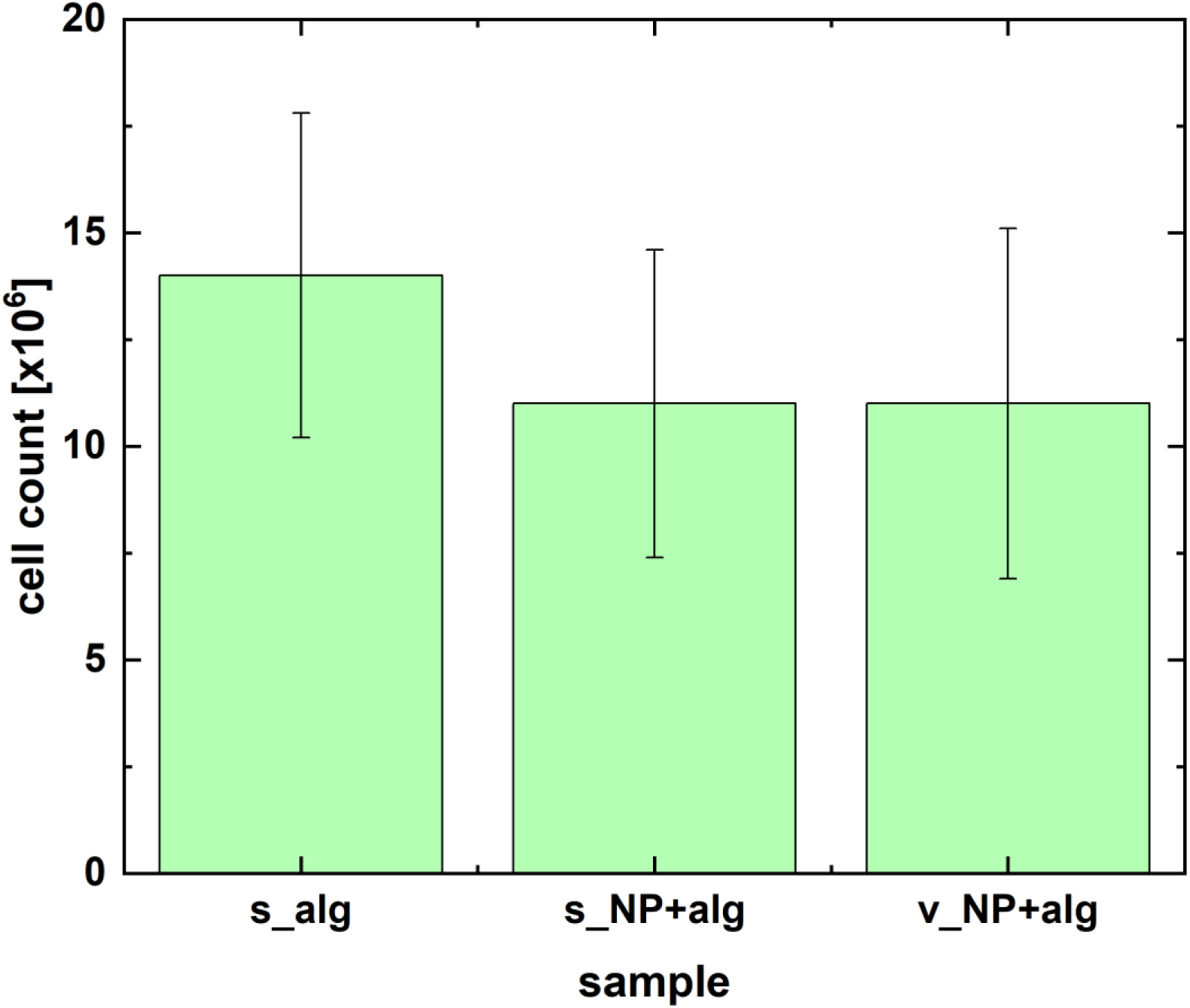
Cell counts of the green microalga (*C. sorokiana*) in bioprinted constructs with different geometry and composition after 8 days incubation under a photon irradiance of 6 µmol photons m^-2^ s^-1^. Here s_alg is a bioprinted slab with algae only, while s_NP+alg and v_NP+alg are bioprinted constructs with NPs and algae in a slab and Vgroove geometry, respectively.

**Figure S8:**
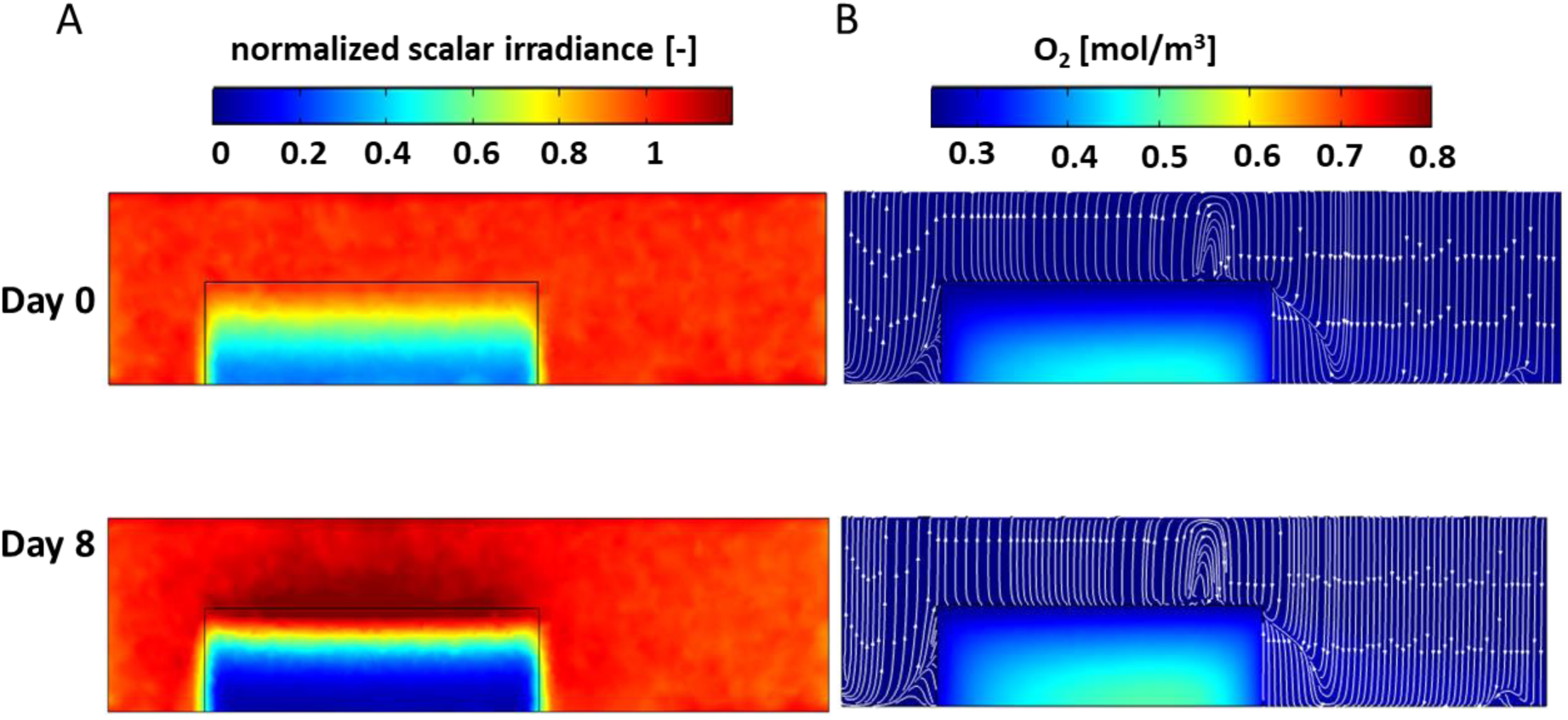
Simulation of light and O_2_ distribution (shown as 2D cross-sections across the peak) in a bioprinted slab with green microalgae in a Vgroove geometry incubated 8 days. A: Day 0 and day 8 light field. B: Day 0 and day 8 O_2_ distribution.

**Figure S9:**
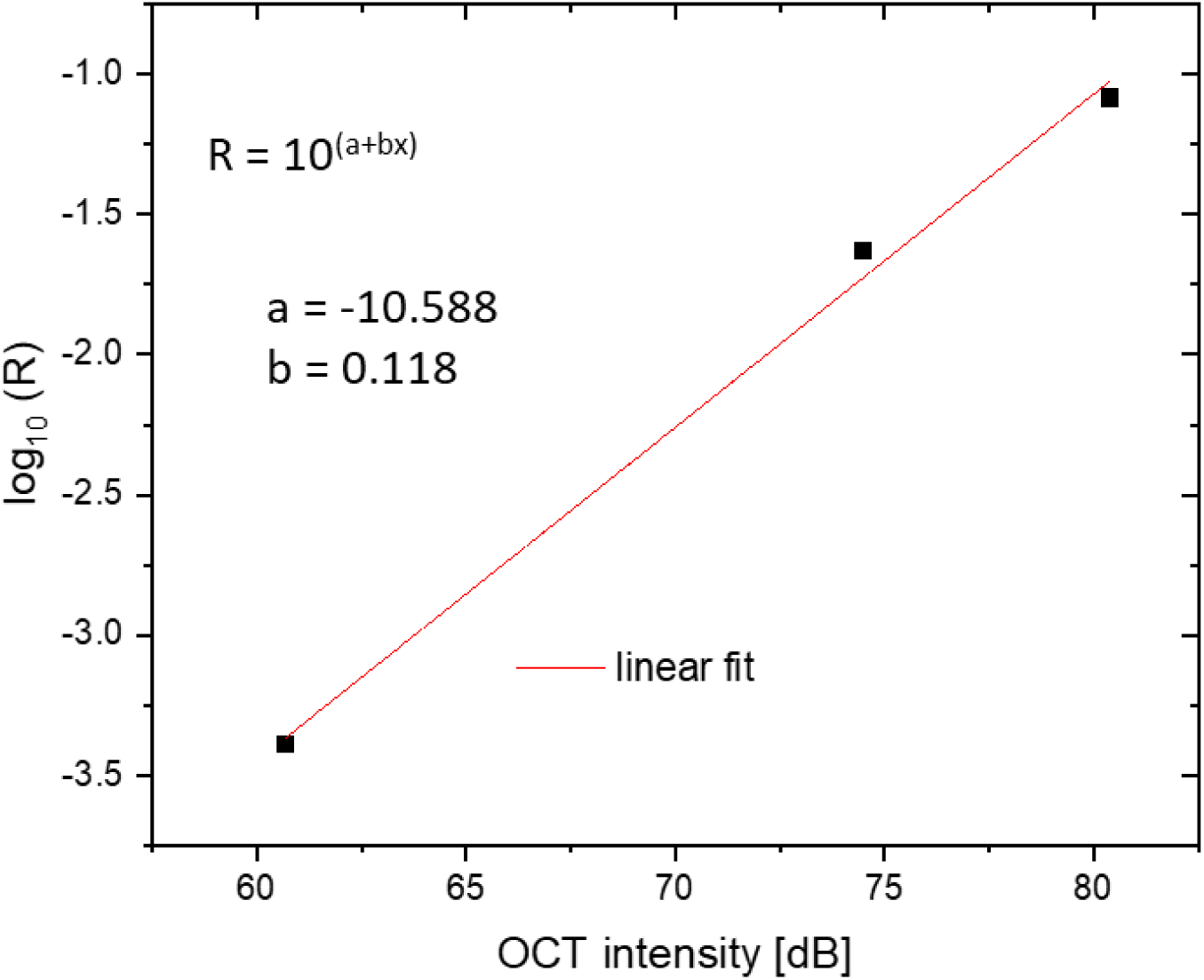
Example of a linear calibration between OCT signal intensity (dB) and reflectivity (log_10_R) data. Three known refractive index mismatches were measured: (1) oil-glass interface, (2) water-glass interface, and (3) air-glass interface.

**Figure S10:**
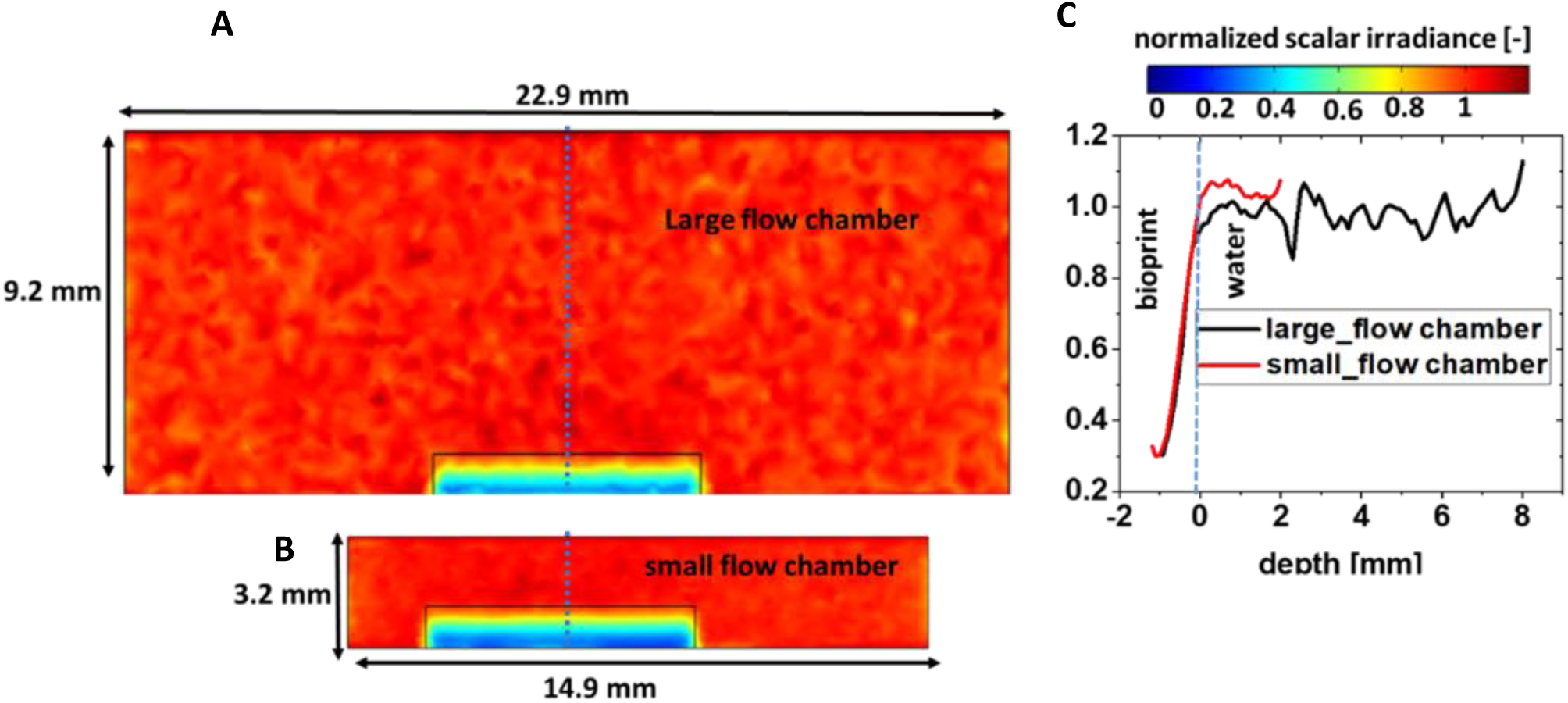
A comparison of MC simulations of the light field (636 nm) in a bioprinted slab geometry when contained in large and small flow chambers using, optical properties from day 8 (Table S8 and Table S10). 2D cross-sectional view of normalized scalar irradiance in **A:** large flow chamber; **B:** small flow chamber. **C:** 1D profiles of light field along the cut lines shown in panels **A** and **B.** The calculated light fields were very similar, with an average normalized scalar irradiance value of 0.61 for large flow chamber and 0.59 for small flow chamber scenario.

**Figure S11:**
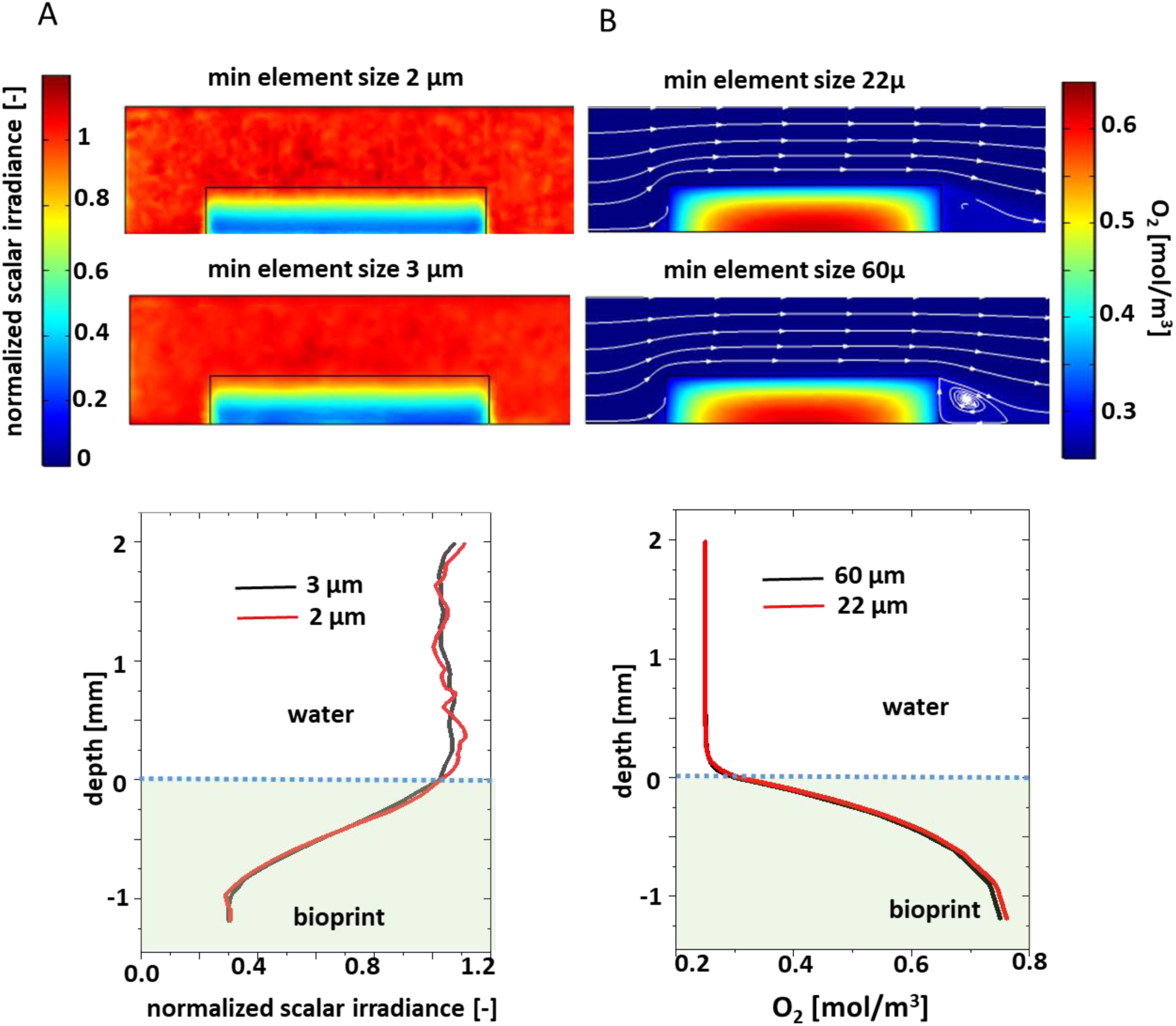
Effects of mesh size on the simulation of light and O_2_ in a bioprinted slab with green microalgae under a photon irradiance (636 nm) of 320 µmol photons m^-2^ s^-1^. Mesh independent study for slab on day 8, **A:** 2D cross-sectional and **C:** 1D plot of normalized scalar irradiance showing that the simulated light field is similar for mesh with a minimum element size of 2 µm and 3 µm. A minimum element mesh size of 3 µm was used for light simulation in the study. **B:** 2D cross-sectional and **D:** 1D plot showing that the simulated O_2_ concentration is almost identical for meshes with a minimum element size of 22 µm and 60 µm. A minimum mesh element size of 60 µm was used for O_2_ mass transfer simulations in the entire study.

**Figure S12:**
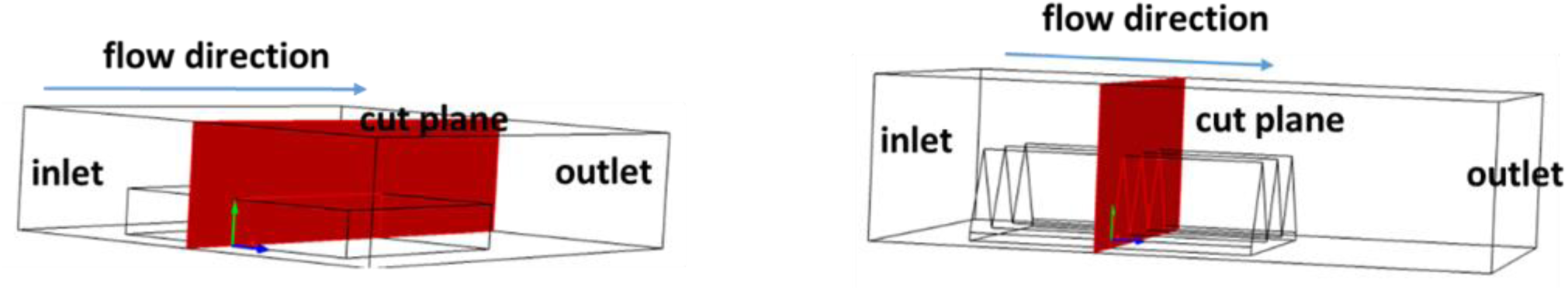
Schematic of the position of the cut-plane (red), in the center of the bioprinted constructs (shown in Fig. 5) with a slab (left) and Vgroove (right) geometry.

**Figure S13:**
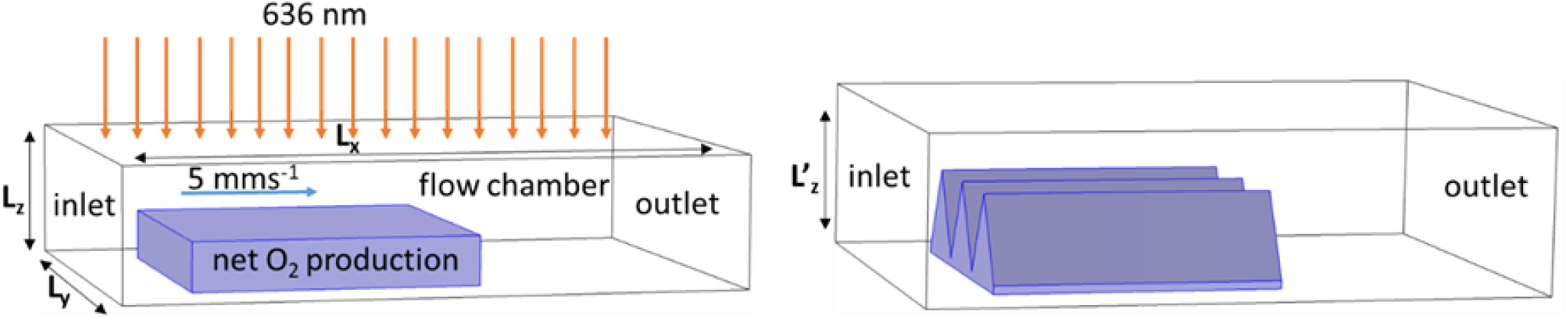
Bioprinted constructs with a slab (left) and Vgroove (right) geometry in a virtual flow chamber used for light, flow and mass transfer simulation (L_x_ 14.92 mm, L_y_ 10.92mm, L_z_ 3.2 mm and L’_z_ 4 mm). A collimated light source (636 nm) illuminates the entire upper boundary of the flow chamber. A no light flux condition was assigned to all other boundaries. The dimensions of the bioprinted constructs are as shown in Fig. 1. Simulations were done with a fully developed laminar flow with an average velocity of 5 mm s^-1^ and a constant O_2_ concentration (100 % air saturation) at the chamber inlet. A fixed gauge pressure of zero and a convection only condition was applied at the chamber outlet. A no slip zero velocity was applied as a boundary condition at the interface between water and bioprinted construct. Symmetry and ‘no flux’ boundary condition were applied on the top and lateral walls of the flow chamber. A net photosynthetic O_2_ production rate, calculated from the MC simulated light field, was assigned to the bioprinted constructs.

**Figure S14:**
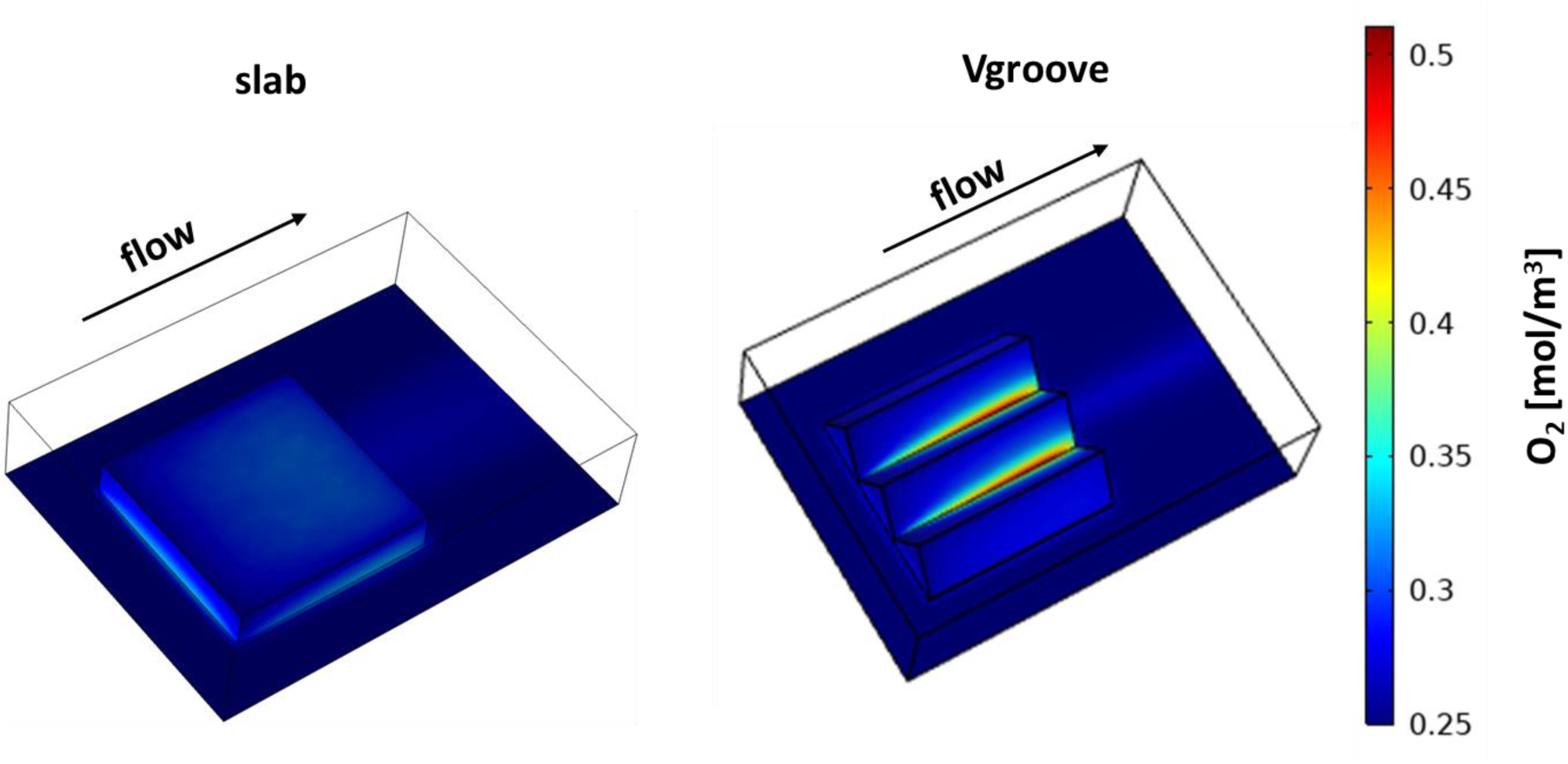
Simulated O_2_ concentrations over bioprinted constructs with a slab (left) and a Vgroove (right) geometry under an incident photon irradiance (636 nm) of 320 µmol photons m^-2^ s^-1^.

